# Temporally specific patterns of neural activity in interconnected corticolimbic structures during reward anticipation

**DOI:** 10.1101/2020.12.17.423162

**Authors:** Megan E. Young, Camille Spencer-Salmon, Clayton Mosher, Sarita Tamang, Kanaka Rajan, Peter H. Rudebeck

## Abstract

Functional neuroimaging studies indicate that interconnected parts of the subcallosal anterior cingulate cortex (ACC), striatum, and amygdala play a fundamental role in affect in health and disease. Yet, while the patterns of neural activity engaged in striatum and amygdala during affective processing are well established, especially during reward anticipation, little is known about subcallosal ACC. Here we recorded neural activity in non-human primate subcallosal ACC and compared this to interconnected parts of basolateral amygdala and rostromedial striatum while macaque monkeys performed reward-based tasks. Applying multiple analysis approaches, we found that neurons in subcallosal ACC and rostromedial striatum preferentially signal anticipated reward using short bursts of activity that form temporally specific patterns. By contrast, basolateral amygdala uses a mixture of both temporally specific and more sustained patterns of activity to signal anticipated reward. Thus, dynamic patterns of neural activity across populations of neurons are engaged in affect, especially in subcallosal ACC.

**HIGHLIGHTS:**

- Sustained changes in neural activity signal anticipated reward in basolateral amygdala
- Temporally specific patterns signal anticipated reward in all areas recorded
- Neurons exhibit more punctate encoding when tasks become more complex
- Temporally specific patterns of neural activity signal different anticipated rewards in BLA

## INTRODUCTION

Corticolimbic areas are central to neural circuit-based accounts of affective processing (Price & Drevets, 2012). Neural activity within human amygdala, striatum, and subcallosal anterior cingulate cortex (subcallosal ACC) is related to processing of affective stimuli and experiences, both rewarding and aversive (Lindquist *et al*., 2012). Damage to or dysfunction within each of these areas is also associated with marked changes in affect (Adolphs *et al*., 1995, 1996).

Of these areas, there has been a particular focus on subcallosal ACC because of its role in pathological changes of affect. Both hyper-and hypoactivity within subcallosal ACC are biomarkers of depression (Mayberg *et al*., 1999; Siegle *et al*., 2012) and activity normalizes following successful pharmacotherapy (Mayberg *et al*., 2000) or cognitive behavioral therapy treatment (Dunlop *et al*., 2017). In schizophrenia, disrupted anticipatory reward responses within subcallosal ACC as well as striatum and amygdala are related to the severity of anhedonia (Dowd & Barch, 2010, 2012). This indicates a specific role for this network of areas in modulating affective responses in anticipation of positive events such as rewards. Consistent with this, lesions (Rudebeck *et al*., 2014) or pharmacological overactivation of subcallosal ACC (Alexander *et al*., 2019) disrupt autonomic arousal and behavior in anticipation of rewards in non-human primates. Thus, in both humans and non-human primates, the circuit linking subcallosal ACC, amygdala and striatum is necessary for affective behavior and specifically responding in anticipation of reward.

Despite this central role for subcallosal ACC in affective behavior, little is known about how neurons within primate subcallosal ACC signal rewards. Unlike striatum and amygdala where the patterns of neural activity related to impending rewards and punishments are well established (Schultz *et al*., 1993; Paton *et al*., 2006), few neurophysiological investigations of primate subcallosal ACC are available. Further, the findings from these studies are equivocal: neurons in subcallosal ACC exhibit almost no encoding of impending reward (Monosov & Hikosaka, 2012), more faithfully encode reward (Azab & Hayden, 2018) or are more strongly modulated by changes between sleep and wakefulness (Gabbott & Rolls, 2013). Notably, a mechanistic account for how subcallosal ACC contributes to reward anticipation is lacking. One possibility is that sustained activity in subcallosal ACC drives arousal through interaction with other areas that directly control bodily states (An *et al*., 1998; Ongur *et al*., 1998). Another possibility is that subcallosal ACC provides a dynamic and temporally specific signal to downstream areas that is then used to sustain arousal in anticipation of reward.

To arbitrate between these two potential alternatives, we took a circuit-level approach and recorded neural activity within subcallosal ACC, basolateral amygdala (BLA), and the part of the striatum where both of these areas project, rostromedial striatum (Haber *et al*., 2006; Cho *et al*., 2013). Recordings were made in two macaque monkeys performing Pavlovian trace-conditioning and instrumental choice tasks for fluid rewards. The Pavlovian trace-conditioning task was adapted from one previously shown to be sensitive to lesions of subcallosal ACC (Rudebeck *et al*., 2014). Here we report that: 1) more neurons in BLA compared to subcallosal ACC or rostromedial striatum signal anticipated reward through sustained activity and 2) individual neurons in subcallosal ACC as well as BLA and rostromedial striatum all exhibit temporally specific patterns of activity in anticipation of reward. Our findings indicate that anticipated reward is signaled in these areas by different encoding schemes and subcallosal ACC primarily engages temporally specific patterns of activity to signal reward.

## RESULTS

### Behavioral and autonomic correlates of anticipated reward

We trained two macaque monkeys (monkeys H and D) on a modified version of the Pavlovian trace-conditioning task previously shown to be sensitive to lesions of subcallosal ACC (**Figure 1A**). First, monkeys were extensively trained to maintain gaze on a centrally presented spot for 2.8 - 3 seconds for three small drops of fluid. Then a Pavlovian trace conditioning procedure was superimposed on this fixation task. On 75% of fixation trials, stimuli of equal luminance associated with subsequent delivery of either juice (CS+^juice^), water (CS+^water^) or nothing (CS-) were presented for one second with equal probability shortly after fixation was acquired. If CS+^juice^ or CS+^water^ stimuli were presented a large drop of the corresponding reward (0.5 ml) was delivered 0.5-0.6 seconds after stimulus offset. Presentations of the CS-were associated with no reward being delivered. In the remaining 25% of trials, no CS was presented. To probe responses to unsignaled rewards, on 20% of these unsignaled trials (5% of total trials), a reward was delivered 2.1-2.7 seconds after fixation onset. These unsignaled reward trials are not included in the following analyses.

**Figure 1:**
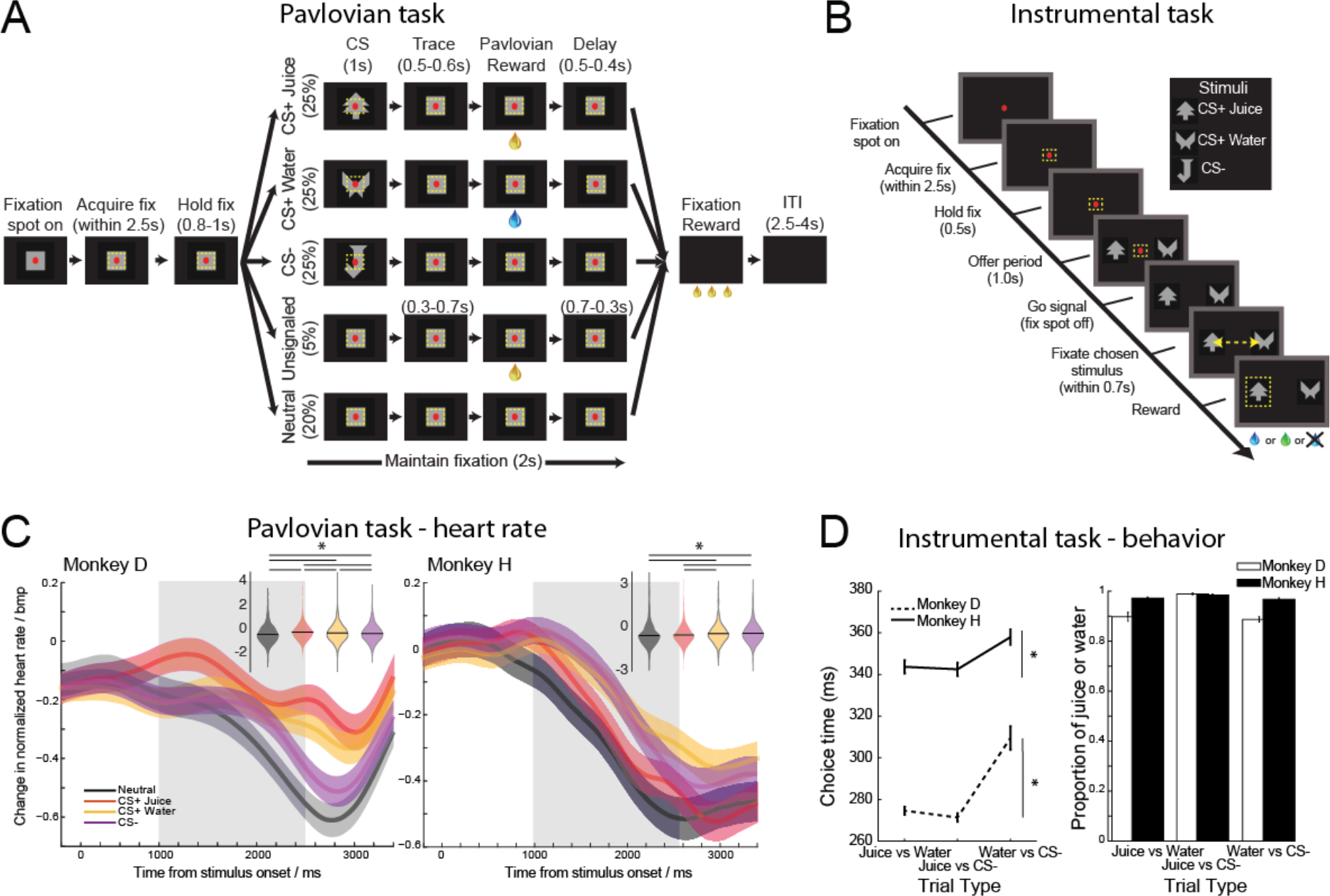
Autonomic and behavioral responses in Pavlovian and instrumental tasks. **A**) In the Pavlovian task, monkeys were required to maintain gaze on a centrally located red spot for 2.8–3.0 seconds to receive three small drops of juice reward. In 75% of trials, one of three conditioned stimuli, either CS+^juice^, CS+^water^ or CS–, were presented for 1 s (CS period) before a 0.4–0.5 second trace interval. If either the CS+^juice^ or CS+^water^ was presented, then 0.5 ml of the corresponding fluid reward, either juice or water, was delivered at the end of the trace interval. **B**) In the instrumental task, monkeys were required to maintain gaze on a centrally located spot. Then two stimuli were presented to the left and right for 1 s. Stimuli were the same as those presented in the Pavlovian task and had the same corresponding reward assignments. The central spot was then extinguished, and this was the cue for monkeys to make their choice by making a saccade to either stimulus. After maintaining fixation on the chosen stimulus for 50 ms the corresponding reward was delivered. **C**) Heart rate as a function of time (percent change, mean ± SEM) for monkeys D and H on neutral, CS+^juice^, CS+^water^ or CS– trials. Inset violin plots show the mean heart rate during the period 1000-2500 ms after stimulus onset (gray shaded region, mean ± SEM). Horizontal lines mark significant pairwise comparisons at p < 0.05. Shaded regions or error bars show SEM. **D**) Choice behavior during the instrumental task. Choice latencies (left) and proportion of choices (right) during the three different trial types. Error bars show SEM.

While subjects performed the task, we monitored both heart rate and pupil size to look for correlates of sustained arousal in anticipation of reward. Both subjects exhibited sustained changes in autonomic arousal after stimulus presentation in anticipation of the fluid rewards that they would receive compared to the neutral condition. In both monkeys, heart rate and pupil size responses were sustained in anticipation of reward that would be delivered (**Figure 1C** and **Supplemental Figure S1**). The pattern of autonomic response differed between heart rate and pupil size measures and was unique to each subject. Heart rate was elevated to both reward-predicting cues in in monkey D compared to CS-or neutral (**Figure 1C**, left, effect of trial, F(3,6480), p<0.0001, pairwise tests, CS+^juice^ > CS+^water^ > CS- > neutral, p<0.05), whereas it was elevated for CS-and CS+^water^ in monkey H compared to both neutral and CS+^juice^ (**Figure 1C**, right, F(3,4258)=12.14; p<0.0001, pairwise tests, CS-or CS+^water^ > CS+^juice^ or neutral, p>0.05). Such variability between subjects is not so surprising given the human literature where autonomic responses to affective stimuli are known to be heterogeneous (Siegel *et al*., 2018). What is critical is that each subjects’ responses were internally consistent and stable over time. In keeping with previous reports (Cash-Padgett *et al*., 2018), pupil size was often constricted in anticipation of receiving juice relative to water and CS-but dilated during the period where monkeys received the reward (**Supplemental Figure 1**).

To confirm that the autonomic responses exhibited by our subjects were related to anticipated reward and that the subjects had learned the CS-outcome relationship, the same two monkeys performed an instrumental choice task (**Figure 1B**). The instrumental task used the same stimulus-reward pairings presented during the Pavlovian task and the two tasks were predominantly run one after the other in each recording session. In the instrumental choice task, both subjects chose CS+^juice^ over either CS+^water^ or CS-on more than 95% of trials (**Fig. 1D**). A similar preference was apparent on trials where they could only choose between CS+^water^ and CS. The choice response latency, the amount of time from the go signal to the selection of one of the stimuli, was also modulated by subjects’ preferences. Monkeys were faster to respond on trials where a CS+^juice^ stimulus was presented (**Figure 1D**, effect of reward, F(1,3)=13.44, p<0.00001). Thus, monkeys exhibited differential patterns of sustained autonomic arousal after stimuli were presented but exhibited choices and response latencies that were similar, consistent, and qualitatively matched the motivational significance of the stimuli.

### Neural activity in subcallosal ACC, BLA, and rostromedial striatum during Pavlovian trace conditioning task

To look for neural correlates of stimulus-linked reward anticipation, we recorded the activity of 656 single neurons and multi-unit responses in subcallosal ACC (n=222), BLA (n=220), and rostromedial striatum (n=214) while monkeys performed the Pavlovian trace-conditioning task. Full details of recordings by area are presented in **Table 1**.

**Table 1:** Counts of neurons and multi-unit responses by area and subject recorded during the Pavlovian-trace conditioning task.

| Area | Monkey H |  |  | Monkey D |  |  | Total |
| --- | --- | --- | --- | --- | --- | --- | --- |
|  | Multi-unit | Single neuron | Total | Multi-unit | Single neuron | Total |  |
| Basolateral amygdala | 20 | 60 | 80 | 26 | 114 | 140 | **220** |
| Subcallosal ACC | 22 | 54 | 76 | 49 | 97 | 146 | **222** |
| Rostromedial striatum | 29 | 69 | 98 | 35 | 81 | 116 | **214** |
| **Total** | **71** | **183** | **254** | **110** | **292** | **402** | **656** |

In all three areas the firing rate of many neurons varied according to the potential for fluid rewards after stimulus presentation. For example, the activity of the subcallosal ACC neuron in **Figure 2A** exhibited maximal firing to the CS+^juice^ and progressively lower firing to CS+^water^ and CS-respectively following stimulus presentation. There was little change in firing rate when conditioned stimuli were absent on neutral trials. Neurons in BLA and rostromedial striatum similarly discriminated between the stimuli presented and the potential reward type (**Figure 2A**, middle and right), but the example neuron in BLA has elevated firing starting just after stimulus onset until reward delivery (middle, **Figure 2A**).

**Figure 2:**
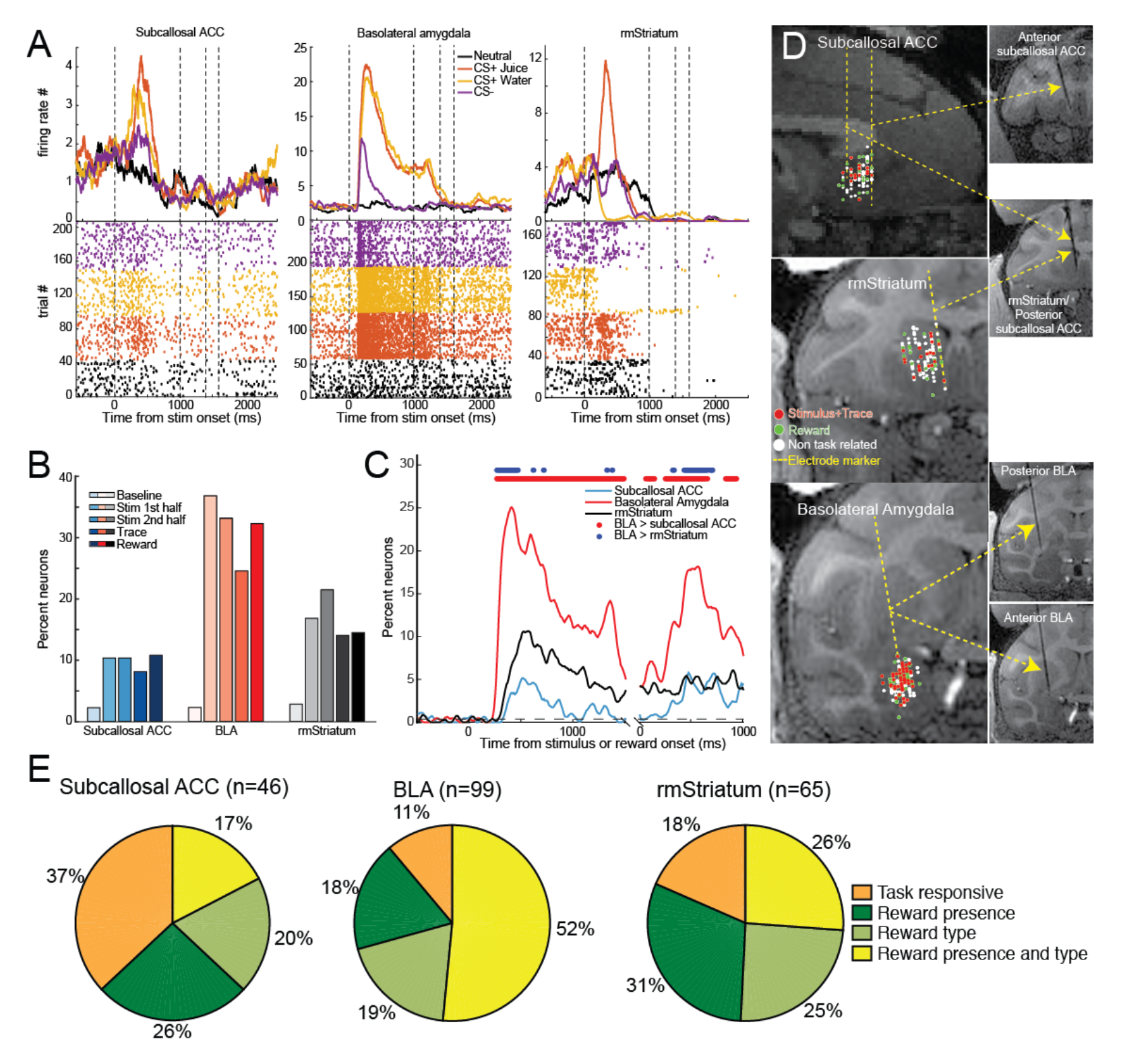
Neural activity in subcallosal ACC, BLA and rostromedial striatum during the Pavlovian task. **A**) Spike density functions and raster plots depicting the activity of example neurons recorded within subcallosal ACC (left), BLA (middle) and rostromedial (rm) striatum (right). Neurons in subcallosal ACC and rostromedial striatum exhibit the highest firing rate to stimuli associated with receiving juice reward and progressively lower firing to the other stimuli. Note that activity returns to baseline before the trace interval in both. The neuron in BLA (middle) exhibits higher and sustained firing to the stimuli associated with juice and water. The color code shows the different trial types. **B**) Percent of neurons in subcallosal ACC, BLA and rostromedial striatum classified by a sliding ANOVA as encoding the different task conditions during either the *baseline period* (0.5 s before the onset of stimulus), the *stimulus period 1^st^ half* (0–0.5 s after the onset of the stimulus), the *stimulus period 2^nd^ half* (0.5–1 s after the onset of stimulus), the *trace interval* (1-1.5s after the onset of the stimulus), or the *reward period* (0-0.5s after reward onset). The percent classified during the baseline period indicates the false discovery rate. **C**) Time course of stimulus–reward encoding in subcallosal ACC (blue), BLA (red) and rostromedial striatum (black) following the presentation of the conditioned stimuli. Red and blue dots at the top indicate significant differences in the proportion of neurons between areas (p<0.0167, Gaussian approximation test with false discovery rate correction). Dotted line depicts the data derived false discovery rate at each timepoint. **D**) Location of recorded neurons in the subcallosal ACC (top, sagittal view), rostromedial striatum (middle, coronal view) and BLA (bottom, coronal view). Insets show the location of electrodes on T1-weighted MRIs targeting each of the three structures. Each dot represents a neuron. Red and green dots denote neurons classified as encoding trial type during the combined stimulus and trace or reward interval, respectively. **E)** Percentage of classified neurons in subcallosal ACC, BLA and rostomedial striatum that also signaled the presence or absence of reward (CS+ versus CS-, green), the reward type (CS+^juice^ versus CS+^water^, red), both (yellow) or were simply task responsive (orange).

To quantify encoding differences between areas we first computed smoothed spike rate functions for each neuron on each trial using a 201 ms window in 10 ms steps on a period of 2.5 seconds starting 500 ms before stimulus presentation and ending 2000 ms after stimulus presentation. These smoothed spike rate functions were then analyzed using an ANOVA producing an array of 250 p-values for each neuron. This array was then divided into five equal epochs of 500 ms/50 bins; a baseline period (−500 – 0 ms), stimulus 1^st^ half (0 – 500 ms), stimulus 2^nd^ half (500 – 1000 ms), trace (1000 – 1500 ms), and reward (0-500 ms after reward delivery). Neurons were classified as encoding a task variable within one of these epochs if they met a threshold of p < 0.007 for 3 consecutive steps within that epoch and this is what is shown in Figure 2B. To provide further resolution on the time course of activation, the proportion of classified neurons at each bin/time point is shown in Figure 2C. The threshold was determined by selecting values that produced a false discovery rate during the baseline period of < 2-3% (see **Experimental Procedures**). Thus, the proportion of neurons classified as encoding trial type during the baseline period provides an estimate of the false discovery rate in **Figures 2B** and **C**. What we describe below is the combination of both single and multiunit responses as there were no differences between analyses conducted on either dataset (**Supplementary Figure S2**). Similarly, data are collapsed across both subjects there were few differences (**Supplemental Figure S3**).

Following presentation of the stimuli, a greater proportion of neurons in BLA encoded the different task conditions than those in either subcallosal ACC or rostromedial striatum (Gaussian approximation test with false discovery rate correction, p<0.0167, **Figure 2B** and **C**). During the trace period when animals had to remember if a reward-predicting stimulus had been presented, more BLA neurons encoded impending reward delivery than in either subcallosal ACC or rostromedial striatum (**Figures 2B** and **C**). Note that in **Figure 2C** the proportion of subcallosal ACC neurons encoding the task conditions was statistically different to BLA in nearly all of the stimulus, trace, and rewards periods although it exceeded chance in all of these periods (compare blue and dashed line). This indicates that when pooled across time, subcallosal ACC neurons encode reward anticipation, but they do not so do in the same sustained manner, as in BLA.

To further characterize how neurons in each of the three areas were signaling different task conditions we conducted an additional set of analyses on neurons classified as encoding the during the task. First, we investigated whether neurons in the three areas predominantly signaled reward conditions by either increasing or decreasing activity during the stimulus period. We found that neurons in both subcallosal ACC (increase/decrease: 30/16) and BLA (increase/decrease: 60/39) were more likely to increase activity in response to the different conditions, whereas the opposite was true in rostromedial striatum (increase/decrease: 27/38). Despite this variation between areas, none of the proportions for the three areas were statistically different from an even split (all three comparisons, X^2^<1.65, p>0.05).

Next, we conducted two additional analyses on neurons classified as encoding during the task using the same sliding ANOVA approach that we detail above. First, we looked for neurons that discriminated cued conditions that would lead to reward as opposed to those that would not (i.e., CS+^juice^ and ^water^ vs CS-and neutral). From here on we refer to neurons classified by this analysis as encoding reward presence. For the second we looked for neurons that discriminated between the different types of reward, (i.e. CS+^juice^ vs CS+^water^). From here on we refer to neurons classified by this analysis as encoding reward type. Note by conducting two independent analyses it is possible for a neuron to be classified as significantly discriminating both reward presence and type. As before we used a threshold of 3 consecutive bins at p<0.007 and confirmed that this produced a false discovery rate of less than 3% in each case.

These additional analyses revealed that the activity of the majority of task responsive neurons in all areas discriminated between the potential for and type of reward in both the stimulus and reward periods of the task (**Figure 2E**). The proportions of neurons encoding reward presence and type varied between the recorded areas. In the stimulus period, more neurons in BLA compared to subcallosal ACC discriminated the potential for reward (subcallosal ACC: 20/46; BLA: 69/99, X^2^=2.83, p<0.005) and different types of reward (subcallosal ACC: 17/46; BLA: 70/99, X^2^=3.68, p<0.001). When we compared encoding in BLA and rostromedial striatum there was only a difference between the proportion of neurons encoding reward type (rostromedial striatum: 33/65; BLA: 70/99, X^2^=2.42, p<0.02) but not reward presence (rostromedial striatum: 37/65; BLA: 69/99, X^2^=1.51, p>0.1). Comparison of the proportion of neurons encoding reward presence or type in subcallosal ACC and rostromedial striatum failed to find any differences between areas (both comparisons, X^2^<1.2, p>0.2). A similar set of analyses conducted on the reward period produced fewer clear distinctions between areas (see Supplemental information, **Figure S3**). In summary, a higher proportion of neurons in BLA compared to subcallosal ACC and rostromedial striatum signaled reward type in the stimulus period, further highlighting the stronger encoding of reward in this area.

To return to addressing our two competing hypotheses concerning the mechanisms engaged during reward anticipation, we next looked at whether encoding was more sustained or dynamic across the three recorded areas. In contrast to neurons in BLA, anticipated reward encoding in subcallosal ACC as indexed by the percent explained variance was less sustained (**Figure 3**, bottom). Indeed, when we quantified the length of anticipated reward encoding in classified neurons across the three areas, BLA neurons exhibited longer encoding in anticipation of reward than either subcallosal ACC or rostromedial striatum (Kruskal-Wallis test on number of significant bins in stimulus and trace period, χ^2^>23.45, p<0.001, BLA versus either subcallosal ACC or rostromedial striatum, χ ^2^>9.36, p<0.005, **Figure 3**, top). There was no difference between the length of encoding in subcallosal ACC and rostromedial striatum (p>0.05).

**Figure 3:**
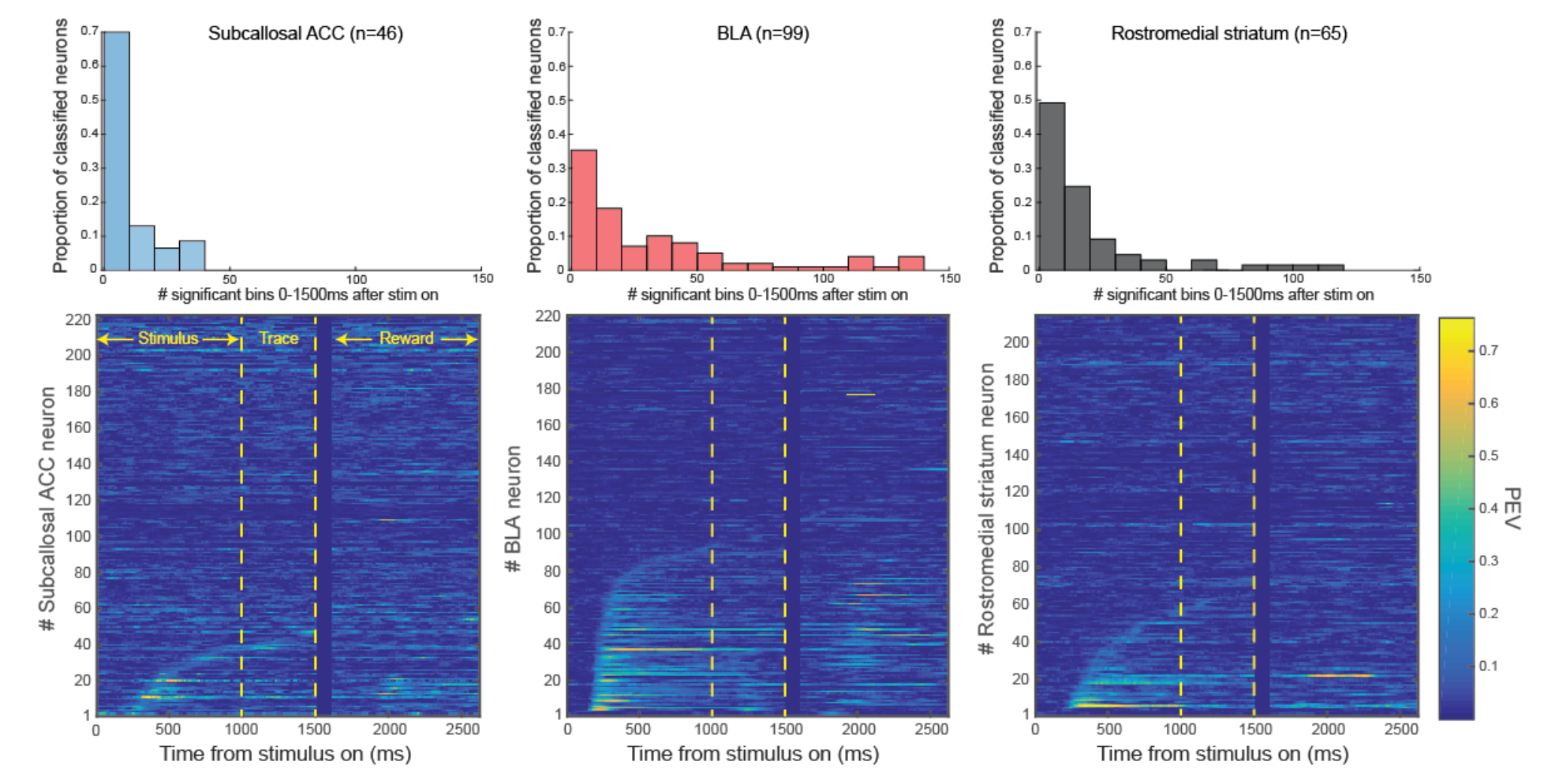
Timing and length of encoding of the different trial types in subcallosal ACC, BLA and rostromedial striatum. Percent explained variance (PEV) associated with the different conditions for each neuron (bottom) and counts of the number of significant bins (top) for subcallosal ACC (left), BLA (middle), and rostromedial striatum (right) relative to stimulus and reward onset. In the plots of PEV (bottom), neurons are sorted according to the first bin in which they were classified as being significantly modulated. Lighter or ‘hotter’ colors are associated with higher explained variance. Because of the variable trace interval, data are temporally realigned for the reward period and there is a period which is intentionally left blank/dark blue between 1500- 1600ms. Histograms (top) of the length of encoding for neurons classified as encoding in each area.

Structural MRI scans with recording electrodes in place (**Figure 2D**) and reconstructions based on these scans confirmed that recording locations were well distributed within the areas targeted. Thus, our data indicate that there are differences in the level of sustained encoding for reward across the three brain areas recorded; BLA exhibits longer encoding than both rostromedial striatum and subcallosal ACC. Taken together, this result appears to argue against the hypothesis that subcallosal ACC uses sustained activity to encode impending reward.

### Neural activity in subcallosal ACC, BLA, and rostromedial striatum during instrumental choice

The absence of sustained encoding in subcallosal ACC in the Pavlovian task could theoretically be because such encoding is only present in settings that require instrumental actions to specific locations to obtain reward. Encoding of both reward and stimulus location in subcallosal ACC during instrumental tasks suggests that action contingency could be a key factor (Strait *et al*., 2016). To address this possibility, we recorded the activity of 421 single neuron and multiunit responses in subcallosal ACC (n=109), BLA (n=144), and rostromedial striatum (n=168) while monkeys performed the instrumental choice task where they chose between pairs of stimuli that had been presented in the Pavlovian trace conditioning task on each trial. A full breakdown of this information is provided in **Table 2**.

**Table 2:**
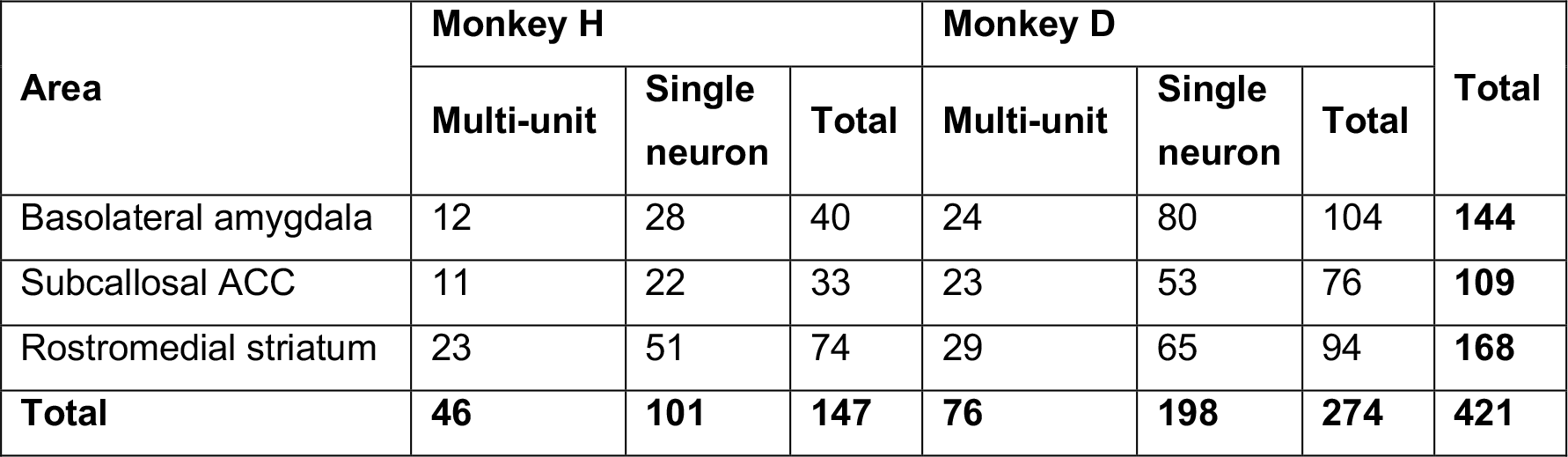
Counts of neurons and multi-unit responses by area and subject recorded during the Instrumental-choice task.

| Area | Monkey H |  |  | Monkey D |  |  | Total |
| --- | --- | --- | --- | --- | --- | --- | --- |
|  | Multi-unit | Single neuron | Total | Multi-unit | Single neuron | Total |  |
| Basolateral amygdala | 12 | 28 | 40 | 24 | 80 | 104 | **144** |
| Subcallosal ACC | 11 | 22 | 33 | 23 | 53 | 76 | **109** |
| Rostromedial striatum | 23 | 51 | 74 | 29 | 65 | 94 | **168** |
| **Total** | **46** | **101** | **147** | **76** | **198** | **274** | **421** |

When monkeys had to choose between options by making an eye movement to obtain a reward, the firing rate of many neurons across the three recorded areas varied with the identity of the chosen reward, either juice or water (**Figure 4A** and **B**). Few neurons encoded the movement direction that monkeys would make or the interaction between chosen reward and movement direction (**Supplementary information, Fig S4**). Additional analyses also confirmed that encoding was best characterized by reward outcome as opposed to aspects of choice difficulty.

Similar to the Pavlovian task, neurons in BLA signaled the chosen reward to a greater extent than both subcallosal ACC and rostromedial striatum (Gaussian approximation test with false discovery rate correction, p<0.0167, **Figure 4C**). Again, the length of encoding within BLA was greater than in both subcallosal ACC and rostromedial striatum, but in the instrumental task this difference did not reach statistical significance (p>0.1, **Supplementary Figure S5**). Indeed, unlike the Pavlovian task there was little sustained encoding of reward in any of the areas. In summary, when monkeys had to make instrumental actions to receive rewards, neurons in subcallosal ACC again did not preferentially use sustained encoding to signal impending reward.

**Figure 4:**
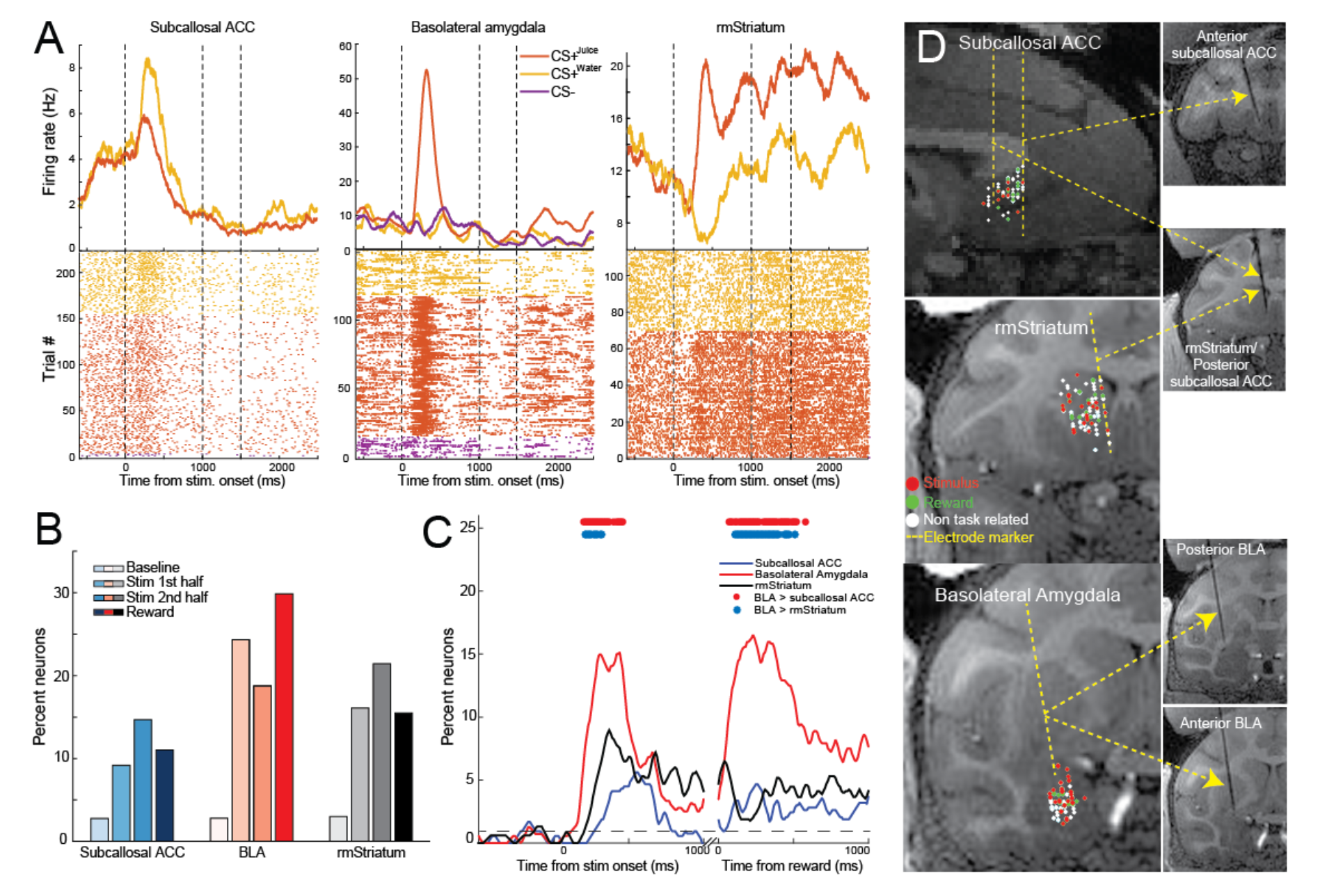
Neural activity in subcallosal ACC, BLA and rostromedial striatum during the instrumental choice task. **A**) Spike density functions and raster plots depicting the activity of example neurons recorded within subcallosal ACC (left), BLA (middle) and rostromedial (rm) striatum (right). The neuron in subcallosal ACC exhibits higher firing when the stimulus associated with water will be chosen. Neurons in rostromedial striatum and BLA exhibit the highest firing rate when juice will be chosen. The color code shows the different trial types. **B**) Percent of neurons in subcallosal ACC, BLA and rostromedial striatum classified by a sliding ANOVA as encoding a the trial types during either the *baseline period* (0.5 s before the onset of the stimuli), the *stimulus period 1^st^ half* (0–0.5 s after the onset of the stimulus), the *stimulus period 2^nd^ half* (0.5–1 s after the onset of the stimulus), or the *reward period* (0-0.5 s after reward onset). The percent classified during the baseline period indicates the false discovery rate **C**) Time course of encoding in subcallosal ACC (blue), BLA (red) and rostromedial striatum (black) following the presentation of the conditioned stimuli. Red and blue dots at the top indicate significant differences in the proportion of neurons between areas (p<0.0167, Gaussian approximation test with false discovery rate correction). Rostromedial striatum and subcallosal ACC comparison not shown as all p values >0.05. Dotted line depicts the data derived false discovery rate at each timepoint. **D**) Location of recorded neurons in the subcallosal ACC (top, sagittal view), rostromedial striatum (middle, coronal view) and BLA (bottom, coronal view). Insets show the location of electrodes on T1- weighted MRIs targeting each of the three structures. Each dot represents a neuron. Red and green dots denote neurons classified as encoding trial type during the stimulus period and reward period, respectively.

### Temporally specific patterns of neural activity in subcallosal ACC, BLA, and rostromedial striatum

If neurons in subcallosal ACC or rostromedial striatum are not using a sustained encoding scheme to signal impending rewards, what is the nature of the mechanism engaged? Mounting evidence indicates that temporally specific patterns of activity within a population of neurons may be a fundamental principle of signaling information in both cortical and subcortical structures (Prut *et al*., 1998; Pastalkova *et al*., 2008; Crowe *et al*., 2010; Harvey *et al*., 2012; Reitich-Stolero & Paz, 2019). Note that we are distinguishing temporally specific patterns of neural activity from sequences, the latter of which have a clear definition based on identification of pattens in simultaneously recorded data. Qualitatively, neurons in subcallosal ACC exhibit temporally punctate bursts of encoding that tile the time interval from stimulus onset to the end of the trace period when visualized and sorted at the population level (**Figure 3**). We thus sought to determine whether these patterns of activity in subcallosal ACC as well as BLA and rostromedial striatum might form temporally specific patterns.

First, to confirm that the patterns of encoding are present in raw firing patterns and not simply in the explained variance, we computed the difference in firing rate between each neuron’s “preferred” and “non-preferred” condition in the Pavlovian task (**Figure 5A**). This approach is analogous to defining a receptive field for each neuron (e.g. Crowe *et al*., 2010). We then sorted neurons in each area by the center of mass (**Figure 5A**). Further aligning these differences in activity by the center of mass revealed that the majority of neurons in subcallosal ACC and rostromedial striatum exhibited punctate changes in activity, indicative of the neurons forming temporally specific patterns of neural activity (**Figure 5A/B**)(Rajan *et al*., 2016). By contrast, neurons in BLA exhibited more sustained changes in firing that were longer in duration than the other two areas (**Figure 5B**, effect of group, F(1,202)=3.27, p<0.05; *post hoc* tests BLA versus ether subcallosal ACC or rostromedial striatum, p<0.05, subcallosal ACC vs rostromedial striatum, p>0.1).

**Figure 5:**
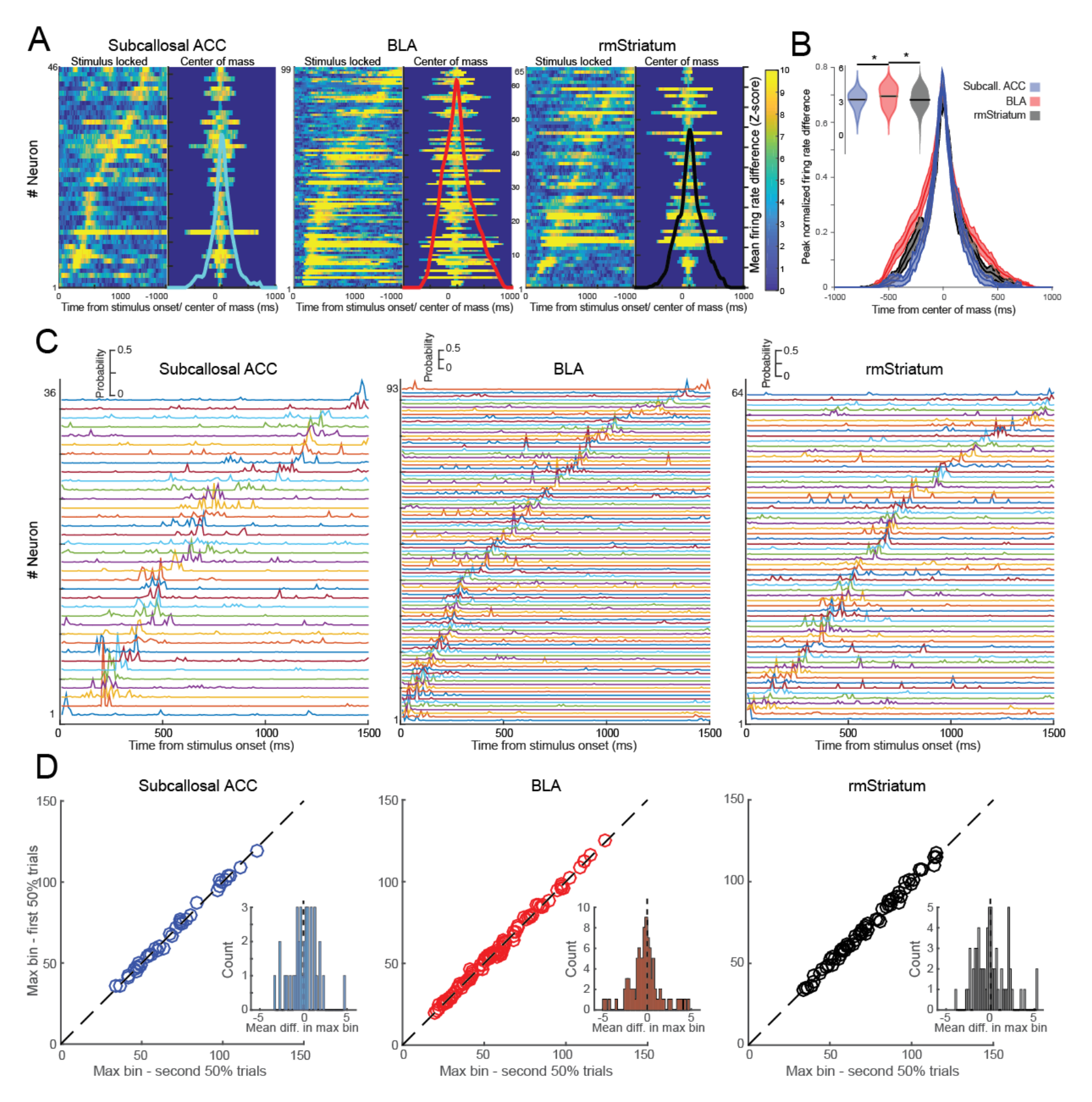
Sequential patterns of neural activity in subcallosal ACC, BLA and rostromedial striatum. **A**) Mean firing rate difference between preferred and anti-preferred trial type for neurons classified as encoding the different trial types in subcallosal ACC, BLA and rostromedial (rm) striatum. Each line represents a single neuron. Firing rate differences are center of mass aligned. Lighter or ‘hotter’ colors are associated with higher differences in firing rate. Solid colored lines are the mean across the population of neurons in each area. **B**) Mean (+/- SEM) peak normalized firing rate difference between preferred and anti-preferred trial types for subcallosal ACC (blue), BLA (red) and rm striatum (black). Inset figure shows log of mean duration of encoding. * denotes p<0.05 **C**) Density of maximum significant encoding times across 500 runs for individual neurons classified as encoding the different trial types in subcallosal ACC (left), BLA (middle), and rostromedial striatum (right). Each colored line represents a single neuron and neurons were sorted according to the time of maximum density. **D**) Mean max bin for first and second 50% of trials for subcallosal ACC (left), BLA (middle) and rm. Striatum. Inset histograms show the distribution of mean difference in bins between first and second 50% of trials.

If neurons in each area form a temporally specific pattern of activity, then individual neurons should have specific times at which they signal impending reward, and this timing should be stable irrespective of the trials that are analyzed. Thus, we next looked at whether neurons across the three areas had specific time points at which they encoded impending reward and how stable/reliable these timepoints were. To ensure statistical power, we selected all neurons that were classified as encoding impending reward in the Pavlovian task that had been recorded for at least 70 completed trials (subcallosal ACC: n=36; BLA: n = 93; rostromedial striatum: n=64). We then randomly sampled fifty percent of the available trials and determined the point of maximal encoding using the previously described sliding ANOVA. The probability distribution of maximal encoding time points for each neuron for 500 such subsamples in subcallosal ACC, BLA and rostromedial striatum is shown in **Figure 5C** sorted by the timing of maximal encoding. Neurons predominantly had specific times at which they encoded different trial types and we confirmed that these maximal encoding time points were statistically different to circularly shuffled data for all neurons (chi-square test on maximal encoding time, p<0.05). Extending this analysis, for each neuron we next compared whether maximum encoding times for one half of the trials drawn at random were different to the other half of trials and repeated this process 500 times. For 96% of neurons there was no difference in maximum encoding times between the two sets of trials (**Figure 5D**, Kruskal-Wallis test, p<0.0167, differences found in subcallosal ACC 2/36=5.71%, BLA 3/93=3.23%, rostromedial striatum 2/64=3.13%). Taken together, these analyses suggest that individual neurons in all areas encoded reward with a robust preference for a particular moment in time prior to reward delivery, a key feature of temporally specific patterns of activity.

To further test the hypothesis that neurons, especially those in subcallosal ACC, are forming temporal temporally-specific patterns of activity to encode reward, we computed a sequential matching index (MI) adapted from (Ji & Wilson, 2007) for each area. This numbering approach can be used to compare two patterns of activity in the same population of neurons, by determining whether each possible pair of neurons appears in the same or opposite order between two patterns of activity or sequences. The resulting matching index, which is equal to the difference between the number of pairs in the same and opposite orders divided by the total number of pairs, is equal to 1 for a perfect match, −1 for a perfect replay of the pattern in opposite order, and 0 for a perfect mixture between same and opposite orderings. Briefly, we subsampled 50% of the trials for each neuron, determined the time of maximal encoding using a sliding ANOVA, and numbered each neuron according to its position in the temporally specific pattern within each area. This temporal ordering was then compared to the same analysis conducted on the other 50% of trials and the MI metric computed. This process was repeated 500 times and compared to the same analysis conducted on randomized data. This analysis revealed that neurons in all three areas exhibited stable ordering that was above chance (**Figure 6A**, subcallosal ACC, MI=0.2196, p<0.035; BLA MI=0.3064, p<0.0004; rostromedial striatum MI=0.2188, p<0.005). These patterns were consistent across monkeys (**Supplemental Figure S6**). Thus, neurons in each of the areas recorded from exhibit stable temporal ordering at the population level, indicative of a temporally specific code.

**Figure 6:**
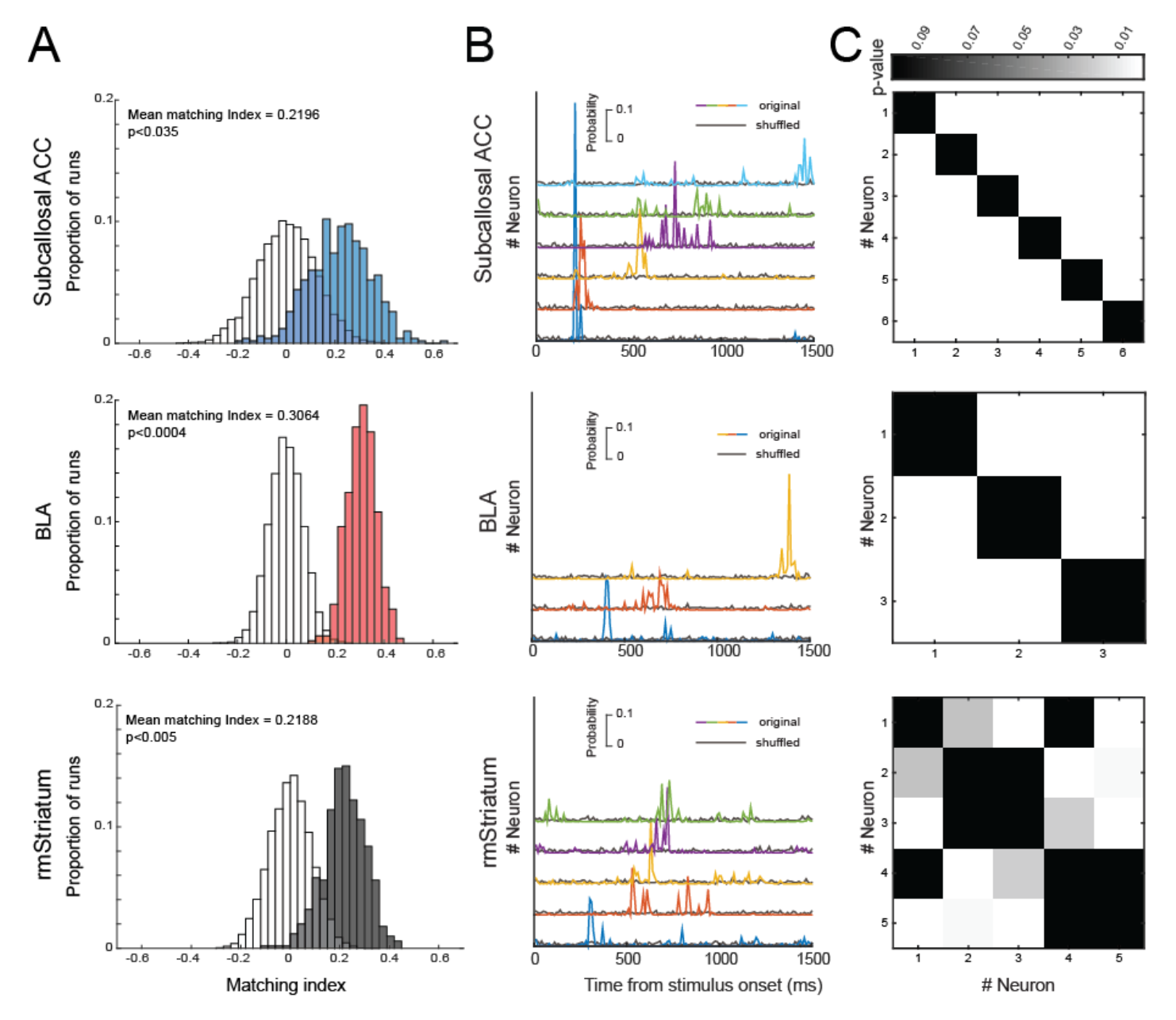
Within session specificity of encoding and changes in encoding length across tasks. **A)** Distribution of matching indices for subcallosal ACC (left), BLA (middle), and rostromedial striatum (right) across the 500 runs. Shuffled data are in grey and actual data are in corresponding colors. **B**) Density of maximum significant encoding times across 500 runs for individual neurons classified as encoding the different trial types in subcallosal ACC (left), BLA (middle), and rostromedial (rm) striatum (right) in a single session. Each colored line represents a single neuron and neurons were sorted according to their maximum density. Black lines represent the density of maximal encoding timepoints for circularly shuffled data. **C**) Statistical comparison of the maximal encoding timepoints for each of the neurons in (B) using a Kruskal-Wallis test. Lighter colors indicate higher level of statistical significance.

Finally, because we recorded neurons across multiple days it is possible that cells in each area encode at a specific time point each day and that we are simply sorting this across-day variability into a temporally specific pattern of neural activity. To address this, we looked for individual recording sessions where more than two neurons in each area were recorded simultaneously. **Figure 6B** shows the diverse time points of maximal encoding of 6, 3 and 5 simultaneously recorded neurons from single recording sessions in subcallosal ACC, BLA and rostromedial striatum, respectively that were classified as encoding reward. Within each area, the maximal encoding time points for each neuron were predominantly different to the other neurons recorded that day (**Figure 6C**, Kruskal-Wallis test, p<0.01). Thus, within all areas recorded neurons exhibited different points of maximal encoding, this was temporally specific, even within a single session, and rarely overlapped with other neurons recorded in the same session. In summary, this series of analyses conducted on neurons classified as encoding the different trial types indicate that neurons in all three recorded areas exhibit patterns of temporally specific neural activity that are stable and reproducible even within a single session.

### Unsupervised identification of temporally specific patterns of neural activity in subcallosal ACC, BLA, and rostromedial striatum

All of the foregone analyses identifying temporally specific patterns of neural activity were conducted on neurons previously classified as encoding the different trial types in the Pavlovian task. If the firing patterns of neurons are truly temporally specific, however, then a greater proportion of neurons should be engaged in these dynamic activity patterns, as it is the timing, not the number of spikes that signal different aspects of the task. Thus, we next investigated whether distinct patterns of neural activity could be identified in pseudopopulations of all of the recorded neurons, not just those classified as encoding the different task conditions. Here we applied an unsupervised computational approach, seqNMF, that has been validated in data from rodents and song birds to extract recurring neural sequences from the activity of populations of neurons (**Figure 7**, Mackevicius *et al*., 2019).

**Figure 7:**
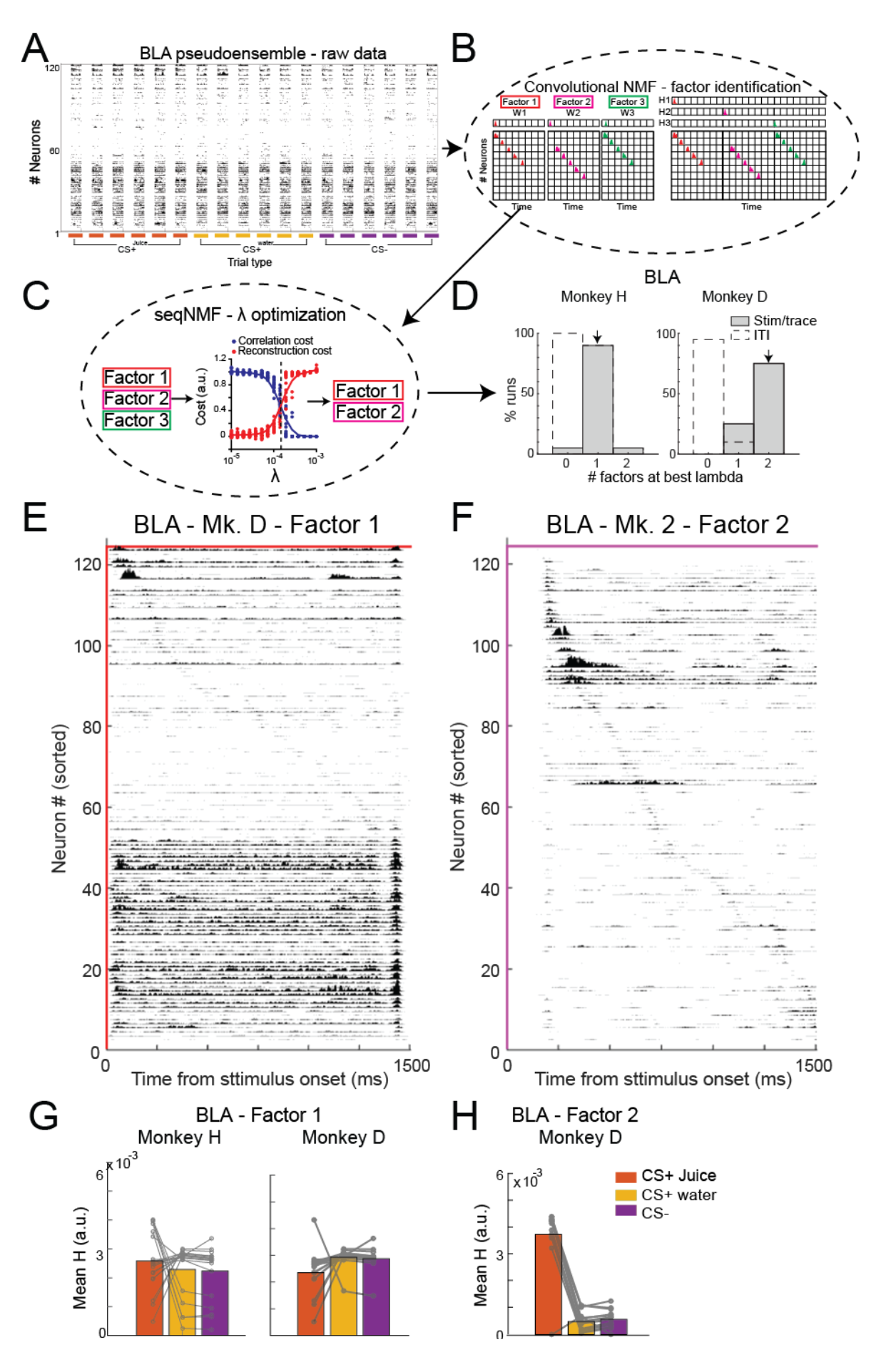
Schematic of factor identification in neural activity data using seqNMF. **A**) An example set of neural activity recorded from 124 neurons in BLA of monkey D (top right). Data are from the 6 trials from each of the three different cued trial types, CS+^juice^ (red), CS+^water^ (yellow), and CS- (purple). **B**) Convolutional NMF models this neural activity data as a neuron by time matrix over the sum of *K* matrices. Matrices constructed for each trial are the product of two components: a non-negative matrix W_k_ of dimension *N* by *L* that stores a sequential pattern of the *N* neurons at *L* time lags. **C**) SeqNMF is a refinement of convolutional NMF as it optimizes a penalty term, λ that removes redundant factors identified by convolutional NMF (Mackevicius et al., 2019). λ is optimized by determining the point at which correlation and reconstruction costs cross (denoted by dotted line where reed and blue curves cross). **D**) When applied to the data recorded in BLA, seqNMF identified one factor in monkey H and two factors in monkey D that correspond to temporally-specific patterns of neural activity. **E/F**) Reconstructions of the two statistically significant factors for monkey D across the 124 recorded neurons. **G/H**) Bar plots of the mean temporal loadings in BLA from significant factors for each monkey. Connected symbols denote temporal loadings for trial types from the individual runs with each factor.

Unlike principal component analysis and clustering, which are typically limited to modeling synchronous activity, seqNMF can model extended spatiotemporal patterns of activity (referred to as factors) and is more scalable than existing techniques aimed at capturing cross-correlations between pairs of neurons. SeqNMF is an extension of convolutional non-negative matrix factorization (convNMF, Smaragdis, 2006) and reduces the probability that a redundant factor will be extracted from the data by adding a cross orthogonality cost term, λ (λ optimization, **Figures 7B** and **C**). Fitting seqNMF to neural activity time-series data outputs n unique factors each represented by a column vector w and their temporal loadings h, a row vector representing the timing and amplitude of those individual factors when they are detected. On each iteration of seqNMF, *h* and *w* are multiplicatively updated by a derivative of the global cost function: *arg min (W, H) = || X - W*H ||^2^_F_ + λ || (W^T^X) SH^T^ || _1, i ∼=j_.* Note that in this case, factors are unique and statistically reproducible patterns of activity. Thus, we used seqNMF to determine whether there were statistically reproducible temporally specific patterns of neural activity in subcallosal ACC, BLA and rostromedial striatum. Further, because this approach as the potential to identify unique temporal loadings and amplitudes, we also looked for differences in the patterns of activity that were distinct between the two subjects.

We applied seqNMF to the firing rates of neurons from 0 to 1500 ms after stimulus onset that had at least 20 instances for each of the CS+^juice^, CS+^water^, and CS-trials. Analyses were conducted separately for each subject to control for individual differences as well as to potentially identify unique patterns of activity in both monkeys. This meant that a total of 115, 124, and 100 neurons in monkey D and 59, 68, and 83 in monkey H from subcallosal ACC, BLA, and rostromedial striatum respectively were combined into pseudopopulations and entered into this analysis. Critically, neurons were not selected based on whether they had previously been classified as encoding the different trial types. Spike trains were smoothed with a 150 ms half Gaussian and parameters were optimized to extract unique and reproducible factors over 20 training and testing runs. Control analyses were conducted on spikes from the last 1500 ms of the ITI and on temporally shuffled or temporally shifted trial segment data using identical parameters to those applied to within trial segments.

Using seqNMF we identified unique factors related to temporally specific patterns of neural activity in all three areas during the stimulus and trace periods of the task (**Figures 7** and **8**). Thus, using an unbiased approach, we found that there are temporally specific patterns of activity in each area, confirming the results of our prior analyses (**Figures 5** & **6**). The amplitude of each factor, as measured by the temporal loadings, and the number of factors identified were, however, different between subcallosal ACC, BLA, and rostromedial striatum and subjects. This indicates that dynamic patterns of activity across the population of neurons signaled different aspects of the task. It also potentially reveals subtle inter-subject differences between the previously identified patterns of temporally specific activity. Importantly, factors were very rarely identified in any of the recorded areas within the ITI (unfilled bars, **Figures 7D** and **8A/B**), when the spike times were randomly shuffled, or when spike activity was circularly shifted in time (**Supplemental Figure S7**). Below we describe the factors identified.

### Subcallosal ACC

A single factor could be reliably discerned from the neurons recorded in subcallosal ACC in both monkeys, and in monkey D where the most neurons were available this factor was closely associated with the anticipation of juice delivery (**Figure 8A, C** and **E** and **Supplemental Figure 8**). **Figure 8C** shows a single factor identified in the activity of neurons in subcallosal ACC from monkey D. Reconstruction of pseudoensemble spiking activity for this factor on three held-out CS+^juice^, CS+^water^, and CS-trials, with the time resolved temporal loadings shown above each trial are shown in (**Supplemental Figure 8A**). In monkey D, this factor had the highest amplitude on CS+^juice^ trials and was equal on CS-and CS+^water^ trials (mean temporal loadings, effect of trial type, F(2,38)=152.1, p<0.0001, *post hoc* tests, CS+^juice^ > CS+^water^ or CS-, p<0.002, **Figure 8E**, right). For monkey H, there was no difference between the temporal loadings for the individual trial types (effect of trial type, F(2,32)=0.74, p>0.48, **Figure 8E**, left).

**Figure 8:**
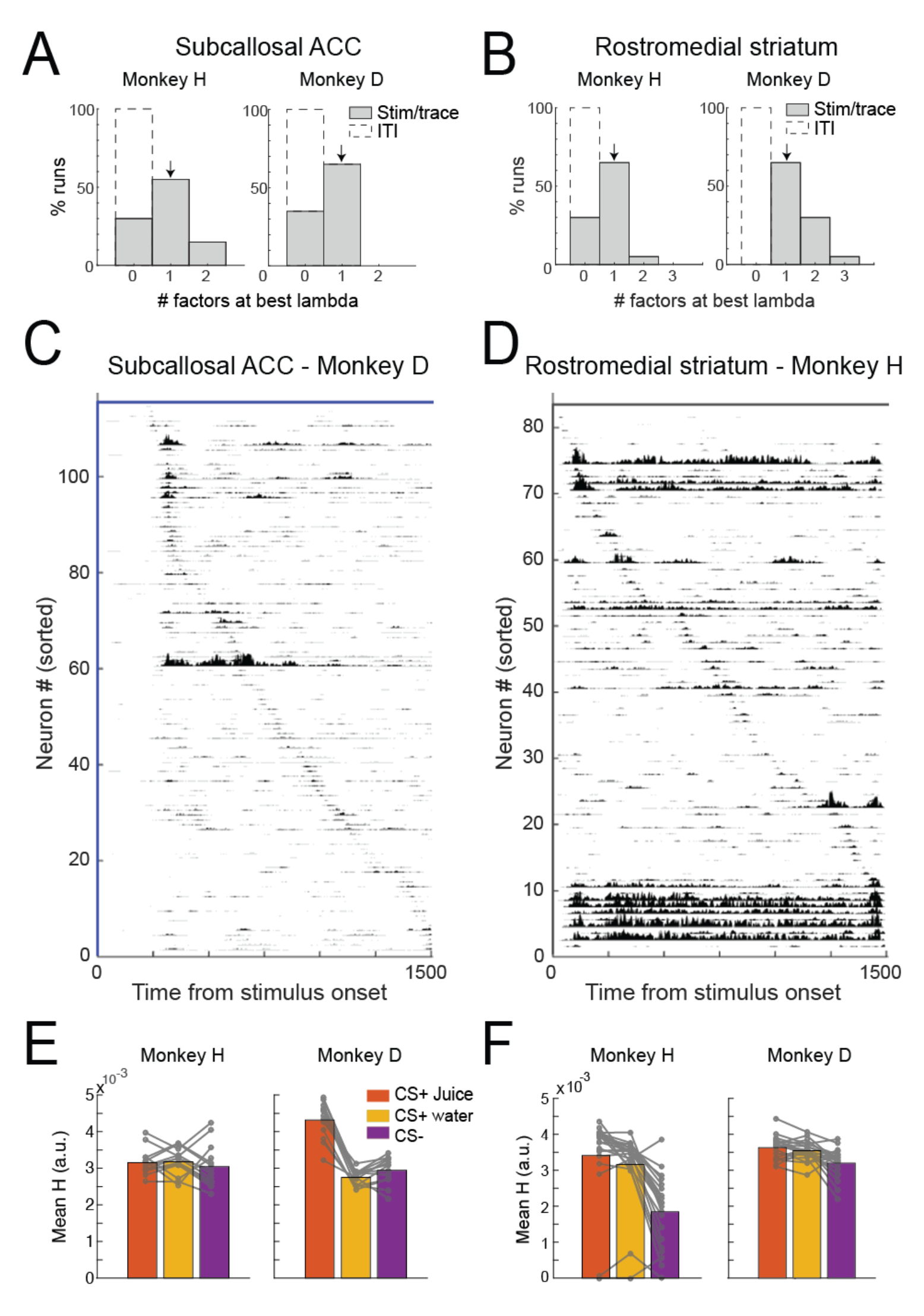
Factors identified in pseudopopulations of subcallosal ACC and rostromedial striatum neurons by seqNMF. **A/B**) Histograms of the number of significant factors identified from the 20 of training and testing runs of seqNMF for monkeys H and D in subcallosal ACC and rostromedial striatum. Factors identified from CS/trace (gray) and ITI (dashed line) area shown. **C/D**) Reconstruction of significant extracted factors from subcallosal ACC (C, monkey D) and rostromedial striatum (D, monkey H). Within each factor neurons are sorted according to the time point of their peak activation. The period shown is from 0 to 1500 ms after stimulus onset which includes both stimulus and trace periods. **E, F**) Bar plots of the mean temporal loadings from significant factors in subcallosal ACC and rostromedial striatum for each monkey. Connected symbols denote temporal loadings for trial types from the runs with factors.

### BLA

Significant factors were identified in the neural activity recorded from BLA in both monkeys (**Figure 7D**). In monkey H, a single factor was found whereas in the neurons recorded from monkey D, two unique factors were identified (**Figures 7D**). The activity of sorted neurons in the two factors identified in monkey D are shown in **Figures 7E** and **F**. As can be seen in the reconstructions of individual trials, the first factor was seen on all trial types and was highly reproducible (**Supplemental Figure 8B**). For this factor the amplitude was equivalent on CS+^juice^, CS+^water^, and CS-trials (effect of trial type, F(2,44)=0.62, p>0.5, **Figure 7G**, right) indicating that it was not associated with the expected outcome (juice, water, or nothing), but instead simply registered the appearance of a stimulus. Reconstruction of the second factor shows that it was predominantly associated with CS+^juice^ trials and was largely absent on CS+^water^ or CS-trials (**Supplemental Figure 8C**, effect of trial type, F(2,44)=145.4, p<0.0001, **Figure 7H**). This is similar to what was seen in subcallosal ACC, where the amplitude of the single temporally specific pattern of neural activity in monkey D was highest on CS+^juice^ trials (**Figure 8C** and **E**).

In the neurons recorded from BLA in monkey H a single factor was most often recovered (**Figures 7D** and **G**, left). This difference between monkeys may be related to the number of neurons available for analysis. Approximately half the number were available from monkey H as compared to monkey D (124 versus 68) and an analysis subsampling the number of neurons in monkey D potentially supports this view (**Supplementary Figure S9**). Unlike in monkey D, the temporal loadings for the factor identified in the BLA of monkey H did not discriminate between the different trial types (effect of trial type, F(2,56)=1.24, p<0.3, **Figures 8E**, right). Note that there were no differences in the proportion of BLA neurons encoding the different trial types between monkeys D and H (**Supplemental Figure S3**). This indicates that it was not simply the case that neurons in monkey H did not discriminate between the different conditions. Thus, individual differences in the factors or the number of neurons available for analysis may explain the differences between animals.

#### Rostromedial striatum

In rostromedial striatum, where equivalent numbers of neurons were available for analysis a single, highly reproducible factor was observed in both monkeys D and H (**Figures 8B, D, and F**). Reconstruction of this factor on spiking data shows that it was closely associated with anticipation of reward delivery (**Supplemental Figure 8D**). Temporal loadings for this factor in both monkeys were higher on rewarded CS+^juice^ and CS+^water^ trials compared to CS-trials (monkeys D and H, effect of trial type, both Fs>8.78, p<0.0001, *post hoc* comparison, CS+^juice^/CS+^water^>CS-, **Figure 8F**). In monkey H, the temporal loadings for this factor not only differentiated between rewarded and unrewarded trials, but also discriminated between CS+^juice^ and CS+^water^ (CS+^juice^ versus CS+^water^, *X*^2^=5.16, p<0.05). This pattern potentially indicates that this factor scaled with the motivational significance of the available rewards, although the same analysis between CS+^water^ and CS+^juice^ in monkey D failed to reach statistical significance (X^2^<1, p>0.3). Thus, the factor identified in rostromedial striatum discriminates rewarded from unrewarded conditions, and in monkey H it also discriminated between all three conditions.

In summary, factors corresponding to temporally specific patterns of neural activity were identified in pseudopopulations of neurons from subcallosal ACC, BLA and rostromedial striatum using unsupervised computational methods. These analyses confirm and extend our previous finding of temporally specific patterns of neural activity and further reveal that: 1) all recorded neurons, not just those classified as encoding the different trial types, contribute to task-specific neural dynamics through the specific timing of their activity, and 2) unique patterns of activity were engaged to signal different aspects of anticipated reward and such patterns were most consistently associated with reward in the rostromedial striatum.

## DISCUSSION

Corticolimbic structures are essential for autonomic and behavioral responses in anticipation of receiving reward. Despite its central role in modulating reward anticipation (Rudebeck *et al*., 2014; Alexander *et al*., 2019), the mechanisms engaged in one part of this network, subcallosal ACC, have been hard to discern. Here we recorded activity within subcallosal ACC as well as interconnected parts of basolateral amygdala and rostromedial striatum in order to gain a circuit-level understanding of how these areas signal reward anticipation. We specifically looked for whether activity in subcallosal ACC either exhibited sustained or more dynamic signals related to anticipated reward.

We found that neurons in BLA exhibited longer, more sustained encoding compared to both subcallosal ACC and rostromedial striatum in anticipation of reward (**Figures 2-4**). By contrast, reward-related activity of single neurons in subcallosal ACC as well as rostromedial striatum was primarily characterized by temporally specific punctate bursts of activity prior to reward delivery. When visualized at the population level, these bursts form temporally specific patterns of neural activity that tile the time until reward is delivered (**Figures 3**, **5** and **6**). Similar patterns of activity were also seen in BLA, but they were intermixed with more sustained encoding (**Figures 3** and **5**).

To further characterize the temporal patterns of neural activity in all of the areas recorded, we first used a previously developed matching index (Ji & Wilson, 2007) to confirm that temporally-specific patterns of task related neural activity were reproducible (**Figure 6**). To further characterize these patterns of neural activity we applied an unsupervised and unbiased computational approach, seqNMF (Mackevicius *et al*., 2019) to our data. During trials where predictive stimuli were presented, seqNMF identified distinct and reproducible factors corresponding to temporally specific patterns in the activity of the pseudopopulations of all neurons recorded from subcallosal ACC, BLA, and rostromedial striatum (**Figures 7** and **8**). In summary, our data reveal that temporally specific patterns of neural activity encode reward anticipation in interconnected cortical and limbic brain areas. Whereas both sustained and temporally specific patterns of encoding were evident in BLA, temporally specific patterns of encoding related to anticipated reward were observed in subcallosal ACC and rostromedial striatum.

### Encoding of anticipated reward in cingulate-amygdala-striatal circuits

Evidence from human neuroimaging studies in healthy individuals (Mayberg *et al*., 1999; Harrison *et al*., 2009), people with psychiatric disorders (Keedwell *et al*., 2005; Dowd & Barch, 2012), as well as anatomical (Ghashghaei *et al*., 2007), and functional studies in non-human primates (Rudebeck *et al*., 2014; Alexander *et al*., 2019) all point to a central role for subcallosal ACC in modulating affect, and specifically reward anticipation. Despite this seemingly central role for subcallosal ACC in affective responses, prior neurophysiology studies looking for signals of anticipated reward in macaque subcallosal ACC have reported virtually no encoding of anticipated reward (Monosov & Hikosaka, 2012), by comparison to amygdala (Paton *et al*., 2006), striatum (Schultz *et al*., 1993), and other parts of medial frontal cortex (Azab & Hayden, 2018).

Our data indicate that one reason that these previous studies may have failed to find strong encoding of impending reward is that they were looking for a population response of neurons time-locked to the presentation of reward-predicting stimuli and/or persistent encoding. Indeed, taking the same analysis approach of aligning the activity of neurons classified as encoding to the onset of predictive stimuli we similarly found that the proportion of neurons in subcallosal ACC and rostromedial striatum signaling impending reward was substantially lower and less sustained in nature than in BLA (**Figures 2-4**). Instead, neurons in subcallosal ACC and rostromedial striatum appear to encode impending reward at specific times before the reward is delivered using punctate bursts of activity. Across recorded neurons, these bursts tile the period until reward is delivered (**Figure 5**) and are specific to a time after stimulus onset (**Figure 6B/C**), a pattern that would have been obscured by standard analytical approaches that average activity across neurons in a population (for example **Figure 2C**).

Recent studies in dorsal ACC and orbitofrontal cortex (OFC) have reported that anticipated reward is signaled using both sustained and temporally specific encoding schemes (Enel *et al*., 2020; Kimmel *et al*., 2020). This is qualitatively similar to what we found in BLA, where both encoding schemes were intermingled (**Figures 2** and **5**). By contrast, in subcallosal ACC as well as rostromedial striatum temporally specific encoding was more prominent. The existence of separable encoding schemes – sustained and temporally specific – is theorized to play different roles in signaling future events. Sustained encoding is postulated to be required to organize behaviors in anticipation of events providing a fixed point for other areas to sample from (Perich & Rajan, 2020). By contrast, temporally specific or sequential encoding is more critical for precisely timing events of interest such as rewards or punishments as well as providing a more efficient way to provide population level-signals to downstream areas (Mello *et al*., 2015). That subcallosal ACC only exhibits one of these schemes further indicates that sustained and temporally specific encoding might be generated by different functional interactions between brain areas, a point we take up below.

Our analyses also revealed other differences between areas related to how single neurons signaled distinct task features. Notably, encoding in subcallosal ACC and rostromedial striatum was less related to the type of reward that would be delivered compared to BLA. While roughly two-thirds of neurons in BLA that were responsive to the task conditions encoded the different types of rewards that would be delivered on each trial, the proportions in subcallosal ACC and rostromedial striatum were substantially lower (**Figure 2F**). This indicates that the encoding of reward in subcallosal ACC and rostromedial striatum appears to be more related to the presence of or the motivational significance of the anticipated reward. Note, that we also found little evidence of spatial position or movement encoding within subcallosal ACC, unlike some previous reports (Strait *et al*., 2016; Azab & Hayden, 2018). A lack of such encoding provides evidence that signals in subcallosal ACC are less indicative of aspects of movement invigoration and are potentially more related to the affective qualities of the anticipated reward.

### Temporally specific patterns of neural activity in anticipation of reward

Sequential patterns of activity across populations of neurons are hypothesized to be an evolutionarily conserved motif for signaling information in neural circuits (Hahnloser *et al*., 2002). While temporally specific sequences of neural activity have been most closely associated with the hippocampus (for example, Pastalkova *et al*., 2008; MacDonald *et al*., 2011) and other medial temporal lobe structures (Reitich-Stolero & Paz, 2019; Vaz *et al*., 2020), task-related sequences of neural activity have also been characterized in other cortical and subcortical areas (Crowe *et al*., 2010; Harvey *et al*., 2012; Mello *et al*., 2015). Because our analyses were primarily at the level of pseudopopulations of neurons recorded on different days, they should not be considered as sequences of neural activity that are generally identified in groups of simultaneously recorded neurons. Our results do, however, provide evidence that sequence-like, temporally specific encoding schemes are present in macaque frontal cortex and are specifically engaged to signal anticipated reward (**Figures 5, 6, 7**, and **8**).

By characterizing temporally specific patterns of neural activity using an unbiased computational approach, seqNMF, we were further able to identify temporal patterns of activity across the whole population of neurons in the three recorded areas, not just those classified as encoding the different task conditions. This not only confirmed our analyses conducted on a more restricted dataset (**Figures 5** and **6**), but also revealed subtle differences in the prominence of the temporally specific patterns of activity depending on the trial conditions or individual subject. In particular, in rostromedial striatum we identified a single unique and highly replicable pattern of activity in both monkeys. Notably, the mean temporal loadings or the amplitude of how discernable the factor was across trial types discriminated between rewarded and non-rewarded trials (**Figure 8**) and in monkey H it discriminated all three conditions (**Figure 8F and Supplemental Figure 8**). Prior work in rodents has identified sequences of neural activity in striatum related to movement and timing behavior (Mello *et al*., 2015). In the Pavlovian trace-conditioning task used here no instrumental actions were required to obtain rewards. Consequently, subjects likely use task events such as stimulus on and offset to predict when rewards will be delivered in time. Thus, our findings indicate that these temporally specific patterns of neural activity in striatum are scaled by reward anticipation.

In both subcallosal ACC and BLA, we also identified temporally specific patterns of neural activity using seqNMF, but here the factors corresponding to unique patterns of activity were less similar between subjects. In subcallosal ACC, a single factor was identified in both subjects, whereas in BLA we identified two distinct factors in monkey D, but only one in monkey H. The amplitude of these factors also differentially scaled between the animals such that in monkey D there was evidence across both subcallosal ACC and BLA that the amplitude of the temporally specific patterns of activity scaled by whether the monkey was anticipating the delivery of juice or not (**Figures 7, 8**, and **Supplemental Figure 8**). Such differential scaling of the patterns of activity was not observed in Monkey H. Note that such differences were only apparent at the level of temporally specific patterns of activity as the proportion of neurons in each subject that signaled different aspects of the task was similar (**Supplementary Figure 3**).

As we noted earlier, such differences in the factors identified by seqNMF in subcallosal ACC and BLA could simply be due to fewer neurons being available for analysis in Monkey H. Alternatively, the divergence in the number of factors and amplitude of the patterns of activity observed in subcallosal ACC and BLA between monkeys D and H potentially provides insight into the individual differences in autonomic responses to the stimuli that were observed in the Pavlovian task (**Figure 1C**). Note that even in large populations of simultaneously recorded neurons differences in the number of factors identified are apparent across subjects (see Figure 8 of Mackevicius *et al*., 2019). A recent study in macaques reported inter-subject variability in behavior and neural activity in the context of information seeking (Jezzini *et al*., 2021) and thus the differences in the patterns of temporally extended activity could be related to individual differences in the task. Confirming the direct relationship between temporally specific patterns of activity in subcallosal ACC, BLA, and autonomic responding will, however, require simultaneous monitoring of neural activity and autonomic activity. This level of resolution was not available here as it necessitates simultaneous recording of large populations of neurons.

### Summary

Our analyses identified that temporally specific patterns of neural activity are engaged to signal anticipated reward, especially in subcallosal ACC and rostromedial striatum. Their presence in both tasks indicates that they are not simply caused by a narrow set of experimental parameters or task settings. Instead, they are likely caused by specific circuit-level interactions. All three areas recorded receive monosynaptic inputs from hippocampus (Barbas & Blatt, 1995; Aggleton *et al*., 2015). As previously noted, neural sequences are prominent in hippocampus during spatial navigation and more recently have been observed in associative learning (Taxidis *et al*., 2020). Cross-talk between hippocampus and subcallosal ACC is essential for adaptive patterns of emotional responding (Wallis *et al*., 2019; Zeredo *et al*., 2019), potentially indicating that interaction with hippocampus could be what drives temporally specific patterns of activity in subcallosal ACC as well as BLA and striatum. Alternatively, temporally specific patterns of activity within a brain area could be the result of local circuit interactions. Either way, determining how this specific neural mechanism contributes to affective regulation in more complex task settings will be essential for determining how cortico-amygdala-striatal circuits contribute to psychiatric disorders characterized by alterations in reward processing.

## EXPERIMENTAL PROCEDURES

### Subjects

Two experimentally naïve adult rhesus macaques (*Macaca mulatta*), one female (Monkey D) and one male (Monkey H), served as subjects. They were 5.6 and 11.0 kg and 16 and 11 years old, respectively, at the beginning of training. Animals were pair housed when possible, kept on a 12- h light-dark cycle, tested during the light part of the day, and had access to food 24 hours a day. Throughout training and testing each monkey’s access to water was controlled for 6 days per week. All procedures were reviewed and approved by the Icahn School of Medicine Animal Care and Use Committee.

### Apparatus

Monkeys were trained to perform Pavlovian and instrumental visually-guided tasks for fluid rewards. Visual stimuli were presented on a 19-inch monitor screen located 56 cm in front of the monkey’s head. During all training and testing sessions monkeys sat in a custom primate chair with their heads restrained. Choices on the screen were indicated by gaze location. Eye position and pupil size were monitored and acquired at 90 frames per second with a camera-based infrared oculometer (PC-60, Arrington Research, Scottsdale, AZ). Heart rate was monitored by recording electrocardiogram (ECG) using a dedicated Biopac recording amplifier (ECG100C Goleta, CA) with the matching IPS100c power supply and MAC110c leads. Fluid rewards were delivered to the monkeys’ mouths using a custom-made pressurized juice delivery system (Mitz, 2005) controlled by a solenoid. All trial events, reward delivery, and timing were controlled using the open source behavioral control software MonkeyLogic (https://monkeylogic.nimh.nih.gov).

### Behavioral task training and testing

Through successive approximation, subjects were trained to maintain their gaze on a centrally-located red spot to receive a fluid reward. The delay to reward was then increased until both subjects were reliably fixating for 4 seconds. A Pavlovian trace conditioning procedure was subsequently superimposed within this fixation task.

Subjects were trained to initiate a Pavlovian trial by fixating on a red fixation point superimposed on a centrally-located gray square for 800-1000 ms. Then, one of four trial types was randomly presented with equal frequency. On CS+^juice^, CS+^water^ and CS-trials, a conditioned stimulus predicting juice (0.5 ml), water (0.5 ml) or no reward was presented behind the fixation spot for 1000 ms in the center of the screen, followed by a 500-600 ms trace interval in which the CS again was replaced by the neutral gray square. After the trace interval, the predicted Pavlovian reward was delivered. On neutral trials, instead of a conditioned stimulus, the neutral gray square remained onscreen after fixation and through the rest of the trial. In one-fifth of such trials, an unsignaled juice reward (0.5 ml) was delivered 1300-1700 ms after fixation. Finally, on all trials, a set of three small (0.1ml) juice rewards was delivered as a reward for successful trial completion 2000 ms after stimulus onset (300-700 ms after Pavlovian reward depending on trial type and trace interval length). Intertrial intervals varied from 2500-4000 ms. Breaking fixation at any point during a trial triggered a timeout interval of 4000 ms before the start of the next trial. Conditioned stimuli varied between subjects and consisted of luminance-matched gray shapes, covering 1.10° of visual angle for monkey D and 2.45° for monkey H.

Subjects also performed an instrumental choice task. Here they made choices between pairs of stimuli from the Pavlovian task (**Figure 1B**). Subjects initiated a trial by fixating on a central red spot for 500 ms, after which two of the three stimuli (drawn by random selection) were shown simultaneously to the left and right side of the fixation spot. All three stimuli were shown with equal frequency with equal probability on each side of the screen. After a variable offer period ranging from 600-2200 ms, the fixation spot turned off, indicating that the monkey had 700 ms to select one of the stimuli by making a saccade and holding fixation on the chosen stimulus for 50 ms. The chosen reward (0.5 ml of juice or water or no reward) was then delivered. Failure to hold fixation when instructed or failure to choose a stimulus within 700 ms of the fixation spot being turned off led to a 4000 ms timeout interval before the next trial. Stimuli for the instrumental task used the same shapes that each subject learned in the Pavlovian task and covered 2.45° of visual angle for both subjects.

### Surgical procedures

All surgical procedures were conducted in a dedicated operating room under aseptic conditions. Anesthesia was induced with ketamine hydrochloride (10 mg/kg, i.m.) and maintained with isoflurane (1.0-3.0%, to effect). Monkeys received isotonic fluids via an intravenous drip. We continuously monitored the animal’s heart rate, respiration rate, blood pressure, expired CO_2_ and body temperature. Monkeys were treated with dexamethasone sodium phosphate (0.4 mg/kg, i.m.) and cefazolin antibiotic (15 mg/kg, i.m.) for one day before and for one week after surgery. At the conclusion of surgery and for two additional days, animals received ketoprofen analgesic (10-15 mg/kg, i.m.); ibuprofen (100 mg) was administered for five additional days.

Each monkey was implanted with a titanium head restraint device and then, in a separate surgery, a plastic recording chamber (27 x 36 mm) was placed over the exposed cranium of the left frontal lobe. The head restraint device and chambers were fixed to the cranium with either titanium screws alone or titanium screws plus a small amount of dental acrylic. Just prior to recording the cranium overlying the recording targets was removed under general anesthesia.

### Physiological and neural recordings

ECG was recorded using surface electrodes (Kendall Medi-Trace 530 Series Foam Electrodes, Covidien, Mansfield, MA) placed on the back of the neck. Placements are varied from day-to-day to avoid irritating a single patch of skin. ECG signals were then low-pass filtered (4 pole Bessel) at 360 Hz, then digitized at 1000 samples/s and recorded on the neurophysiology recording amplifier along with pupil size measurements. Both ECG and pupil size were then processed to remove artifacts (periods of high noise where R-peaks could not be discerned or blinks) using previously validated methods (Mitz *et al*., 2017). After processing all ECG and pupil signals were visually inspected for quality and sessions where there were prolonged periods of noise, signal drop out, or amplifier saturation were excluded from further analysis. This screening process meant that there were 78 and 66 sessions for monkeys H and D available for analysis, respectively.

Potentials from single neurons were isolated with tungsten microelectrodes (FHC, Inc. or Alpha Omega, 0.5-1.5 M at 1 KHz) or 16-channel multi-contact linear arrays (Neuronexus Vector array) advanced by an 8-channel micromanipulator (NAN instruments, Nazareth, Israel) attached to the recording chamber. Spikes from putative single neurons were isolated online using a Plexon Multichannel Acquisition Processor and later verified with Plexon Offline Sorter on the basis of principal-component analysis, visually differentiated waveforms, and interspike intervals. If the waveforms on a channel failed these criteria for single neurons but were differentiated from the noise they were saved and analyzed as multi-unit activity.

Subcallosal ACC recordings were made on the medial surface of the brain ventral to the corpus callosum (**Figure 2D**). All recordings in subcallosal ACC were between the anterior tip of the corpus callosum and the point where structural MRIs were unable to distinguish the medial cortex from the striatum, corresponding to roughly the most anterior point of the septum. This corresponds to between 31.5 to 26.5 mm anterior to the interaural plane. Neurons in basolateral amygdala were primarily recorded in basal and lateral nuclei at least 4 mm ventral to the first neurons that could be discerned after crossing the anterior commissure. Neurons in amygdala were recorded between 22 and 18.5 mm anterior to the interaural plane. Recordings in striatum were made in the rostro-medial segment corresponding to the zone where subcallosal and basal amygdala projections overlap (Haber *et al*., 2006; Cho *et al*., 2013). Neurons in striatum were recorded between 28 to 24 mm anterior to the interaural plane. Recording sites were verified by T1-weighted MRI imaging of electrodes at selected locations in subcallosal ACC, basolateral amygdala and striatum after recordings had been completed (**Figure 2D**).

Neurons were initially isolated before monkeys were engaged in any task. However, in some cases neurons were isolated during the Pavlovian task and these neurons were then recorded in the instrumental version of the task. Other than the quality of isolation, there were no selection criteria for neurons.

### Data analysis

Autonomic and behavioral analysis: To control for across session difference in baseline pupil size and heart rate, data for each session were processed using custom software to remove artifacts and blinks (Mitz *et al*., 2017) and then normalized using a z-score prior to analyses (z = (x-μ) / σ). Pupil dilation was baseline corrected to a 500-ms period extending 250 ms before to 250ms after the onset of CS. This baseline procedure meant that pupil size had the maximum time to stabilize before the presentation of conditioned stimuli (earliest responses are typically after 250ms). Autonomic data were analyzed separately for each monkey using mixed-effects ANOVA models with trial type as a main effect and session as a random effect. For the heart rate analysis, a period from 1800 - 2800 ms after stimulus onset was analyzed. This took into account the time it takes for the heart rate to start changing in response to an external event. For pupil size, a period of 1250- 1750 ms after stimulus onset was analyzed. This took into account both the time for pupil size to start changing in response to an external event as well as the time for the pupil to fully stabilize which is ∼250 ms. Only sessions where there were at least 15 trials of each of the neutral, CS+^juice^, CS+^water^ and CS-were analyzed.

For the analysis of instrumental choices and response times, choices were registered when monkeys made a saccade to one of the two stimuli presented on the screen. For each session, the mean number of choices for each condition was computed and analyzed. Choice response latencies for each trial type were computed by taking the amount of time from the go signal to the selection of one of the stimuli (**Figure 1**). Latencies for each trial type were averaged across sessions and analyzed separately for each monkey using a mixed effects ANOVA with trial type as a main effect and session as a random effect.

Neurophysiology analysis: For analysis of neurophysiology data, only neurons or multi-unit activity that were stably recorded for at least 35 trials were analyzed and it is the activity of those neurons that we report here. Task conditions with too few trials for reliable analysis were not included. Specifically, for the Pavlovian task, the unexpected reward condition was not analyzed as these made up only 5% of total trials. In the instrumental choice task, trials where the CS-was chosen were not included in the analysis as 180/421 units had no CS-choice trials. Overall, there was only a mean of 2.18 trials with CS-chosen per session.

Task-related neurons were identified by fitting a sliding ANOVA model to the firing rates of each neuron in 201 ms bins advancing in 10 ms increments across spike trains recorded in each trial. The ANOVA model included a single parameter of trial type (4 levels) for the Pavlovian task, while the instrumental task was analyzed with a model including chosen stimulus (2 levels, water or juice), response (2 levels, left or right side), and their interaction. Neurons were considered to have significant encoding if the sliding ANOVA returned a p<0.007 for 3 consecutive time bins. These significance criteria were chosen to keep false classification close to 2% or lower for all ANOVA analyses. In the Pavlovian task the sliding window analysis was conducted on 5 equal periods of 500 ms: a baseline period (−500 – 0 ms before stimulus onset), stimulus 1^st^ half (0 – 500 ms after stimulus onset), stimulus 2^nd^ half (500 – 1000 ms after stimulus onset), trace (1000 – 1500 ms after stimulus offset), and reward (0-500 ms after reward delivery). In the instrumental task, the sliding window analysis was conducted on 4 equal periods of 500 ms: baseline (−500 – 0 ms before stimulus onset), stimulus 1^st^ half (0 – 500 ms after stimulus onset), stimulus 2^nd^ half (500 – 1000 ms after stimulus onset), and reward (0-500 ms after reward delivery). Follow up sliding-window ANOVA analyses comparing reward presence (CS+^water^ or CS+^juice^ versus CS-or neutral, 2 levels) and reward type (CS+^water^ or CS+^juice^, 2 levels) during the stimulus and reward periods used the same threshold for classification as above, 3 consecutive bins at p<0.007.

To compare between the proportion of neurons classified in each area we used the Gaussian approximation test on proportions. The threshold for significance, p=0.0167 was adjusted using a false discovery rate correction procedure for time-series data (Hochberg & Benjamini, 1990). Duration of encoding of conditioned stimuli in each area was compared using a Kruskal-Wallis test on the number of significant bins/amount of time of encoding during the combined stimulus and trace period in the Pavlovian task and the stimulus period in the instrumental task. For this analysis the longest contiguous number of bins/amount of time for each neuron was analyzed.

To compare the duration and shape of the patterns of neuronal responses to conditioned stimuli in subcallosal ACC, BLA and rostromedial striatum, we first established the “preferred” and “anti-preferred” conditions for each neuron using the change in z-scored firing rate from baseline after stimulus presentation. First, we calculated a single mean firing rate for each neuron for each condition during the stimulus on (instrumental) or stimulus on and trace (Pavlovian) periods. Then we determined which condition had the maximum (preferred) firing rate and which condition had the minimum (anti-preferred) firing rate for each neuron. Second, we calculated the difference in firing rates for each neuron between the preferred and anti-preferred conditions. Taking the mean firing rates from each condition across trials for 10ms bins (as above), we Z-scored the mean firing rates for the preferred trials and the anti-preferred trials to the baseline activity 500ms before stimulus on. Then the sequence of Z-scored mean firing rates for the anti-preferred condition were subtracted from the Z-scored mean firing rates for the preferred condition to obtain a sequence of preferred vs. anti-preferred firing rate differences for each neuron. Third, we identified the longest segment of differences in Z-scored mean firing rates that was greater than a threshold of 3. These segments were aligned around their centers of mass, peak normalized, and averaged for each area to get a mean response shape for each area. Finally, an ANOVA was conducted on the log normalized length of firing rate differences to compare between brain areas.

In order to assess the stability of peak encoding times for each neuron, a 50% subsample of trials was taken randomly for each neuron 500 times and an ANOVA run on each subsample as above. The time bin with the highest percent explained variance was recorded for each subsample. This analysis included only neurons with significant reward encoding in the entire trial set that were stably recorded for a minimum of 70 trials, to ensure a minimum of 35 trials in each subsample. As a measure of the stability of this peak encoding point, we compared the distribution of maximum encoding points for all sampled 50 percent of trials to that of the remaining 50 percent for all 500 samples for each neuron. We also compared the maximal encoding timepoints between circularly shuffled and unshuffled data taking the same approach as above. In both cases the distributions of maximum encoding points were compared using a Kolmogorov-Smirnov test for each neuron.

To further test whether neurons in each area are encoding reward in a stable temporally specific pattern relative to other neurons in the pseudopopulations, we adapted a form of the matching index analysis developed by (Ji & Wilson, 2007). Here we generated a list of peak encoding times for each neuron by conducting the sliding ANOVA analysis on 50% of the trials for each neuron. Then each neuron was ordered based on these peak encoding times in each area. The same was done for each neuron for the other 50% of trials. This produced two sets of ordered neurons for each area. We then compared the position of each neuron in these two sets of ordered neurons using a matching index that compares the positions of each neuron in the ordered set. The matching index MI is given as MI = (m - n) / (m + n) where m is the number of pairs that are in the same order and n is the number of pairs of neurons that were in a different order. This procedure was repeated 500 times producing a distribution of matching indices. The overall matching index for an area is given as the mean of the matching indices from the 500 repeats. In addition, to confirm that results from the matching index analysis were not simply related to neuron population size, we repeated the above matching index analysis using subsamples of 35 neurons from BLA and rostromedial striatum, the size of the smallest population (subcallosal ACC). Here subsampling the neurons in BLA and rostromedial striatum pseudopopulations did not alter the results.

To assess the significance of the computed matching indices, we calculated the distribution of indices that would be calculated from randomized data to test whether the order of peak encoding times is randomly arranged. To do this, we generated a set of random numbers the length of the number of neurons analyzed in each area and calculated the matching indices for 1,000,000 random iterations. The p-value for each experimental matching index was then calculated as the probability that the matching index would be generated from randomized data of that length. SeqNMF: We characterized the temporally specific patterns in the population neural activity recorded from subcallosal ACC, rostromedial striatum and BLA data using an unsupervised machine learning approach that applies non-negative matrix factorization (NMF) to neural data, seqNMF (Mackevicius *et al*., 2019). seqNMF was developed to extract repeatable patterns of activity from high dimensional data. While the original paper emphasizes the potential applications of seqNMF to simultaneously recorded neurons, this method is agnostic to data type and was developed and tested not only on neural data but also on song spectrograms. For this analysis, we used spiking activity on trials where stimuli were presented (CS+^juice^, CS+^water^ and CS-) starting from when the stimuli were presented to the end of the trace intervals in the Pavlovian task (1500ms total). We included all neurons, irrespective of whether they had been classified as encoding the trial types in the Pavlovian task, that had at least 20 trials for each of the CS+^juice^, CS+^water^, and CS-trials (60 trials total). This yielded 115, 100, and 124 total neurons from subcallosal ACC, rostromedial striatum, and BLA respectively in subject D, and 59, 83, and 68 respectively in subject H. Zero padding was added between trial segments to prevent detection of spurious factors across trial segment boundaries. Neural activity was smoothed with a 150 ms half-Gaussian window. Smoothing with a 50 ms half-Gaussian was also conducted, yielding largely similar results.

SeqNMF differs from other forms of non-negative matrix factorization, such as convolutional NMF (Smaragdis, 2006) in its inclusion of a penalty term in the overall cost function called the x-ortho penalty, the magnitude of which is scaled by the hyperparameter, λ. The x-ortho penalty suppresses the extraction of multiple factors to explain different instances of the same factor and is computed as such: for a given factor in W, we compute its overlap with the data at time t (*W^T^X*). We then compute the pairwise correlation between the smoothed temporal loading (SH) of each factor and the overlap of every other factor in W with the data, such the correlation cost can be written as *λ ||* (*W^T^X*) SH*^T^ ||* _1, i ∼=j_. Because redundant factor will have a high degree of overlap with the data at the same times despite having segregated temporal loadings, this penalty causes any factor that highly overlaps with the data at a given time to suppress the temporal loadings of any other factor that also highly overlap with the data at that time. The overall cost function also incorporates reconstruction cost, or the element-by-element sum of all squared errors between a reconstruction X and original data matrix *X*. The reconstruction X is obtained from the sum of the outer product of w and h, where w is the column vector representing neural factor and h is the row vector representing times and amplitudes at which that factor occurs, such that reconstruction cost = *|| X - WH ||^2^_F_*. Higher values of λ suppress all but one factor to zero amplitude, leading to large reconstruction error if more than one ground-truth sequence exists, while an excessively low λ may cause seqNMF to produce multiple factors to explain different instances of the same factor. As a result, determining the true number of ground-truth temporally specific patterns of neural activity depends on appropriate selection of λ. We implemented the method described in Mackevicius et al., (2019) to select an appropriate λ: we ran 20 iterations of seqMNF at each of a range of values of λ, obtained the average reconstruction error and correlation cost terms at each λ, and chose λ near the crossover point λ_0_ which yielded an approximate balance between the two, either λ =1 λ_0_, or λ =1.5 λ_0_. In the event that most runs returned a single factor, we also compared our factors extracted from optimized seqNMF with factors extracted by setting λ=0 (i.e. non-penalized convolutional NMF). If a similar result was obtained with λ=0, this value was used. As a result of this optimization procedure, λ was set to 0, 0, and 1 for subcallosal ACC, rostromedial striatum, and BLA, respectively.

Once we had chosen an appropriate λ for each area, we re-ran seqNMF for 20 iterations at the selected λ. In order to assess the significance of extracted factors, we employed a 75%/25% training/testing split balanced to have an equal number of trials for each of the three trial types for both the training set and testing set. As a result, seqNMF was trained on 45 trials and tested on 15 trials for each region. For each iteration, we test for factor significance using the held-out test dataset by comparing the skewness of the distribution of overlaps (W’X, where W = factor tensor and X = data) between each factor and the held-out data with the null case. The overlap of a factor with data will be high at timepoints during which the factors occurs, leading to a high skewness of the W’X distribution. From these 20 iterations, we also obtained 1) the temporal loadings for each factor across the time epoch of interest, 2) an average value for reconstruction power (percent variance in the data explained by a given factorization), and 3) the distribution for the number of significant factors found. To compare within and between each factor for each trial type, we conducted mixed-effect ANOVA on the mean of the temporal loadings (H) across the runs for each factor separately with trial type as a main effect and run as a random effect.

Control analyses for seqNMF were conducted using identical free parameters to those optimized for the stimulus and trace interval. First, seqNMF was conducted on the last 1500 ms of the ITI of each trial. This analysis thus maintained the trial progression and spiking statistics but lacked the trial structure. Second, to maintain the trial events in the data, we randomly shuffled the spike times of all neurons on all trials, then smoothed the data, and reran seqNMF. This analysis therefore maintained the overall spike rate in the window of analysis but altered the timing of spikes. Third, to specifically test whether the timing of bursts of spike activity was essential for temporally specific sequences we circularly shifted the neural activity in each trial for each neuron, then smoothed the data, and ran the seqNMF analyses. For example, if on a trial the spike times were all shifted forward by 518 ms, those spikes occurring after 982 ms were moved to the start of the trial in a circular manner. This procedure thus maintained the exact same firing patterns (i.e. inter-spike intervals were maintained) on each trial, but randomly shifted their timing. For each of these control analyses we compared the number of significantly extracted factors to those recovered from non-shuffled data during the stimulus and trace intervals (**Figures 8**, **S7**, and **S8**).

In order to determine whether differences in the number of temporally specific patterns of neural activity identified in the BLA of monkeys H and D was a function of the number of neurons available for analysis, we conducted a subsample analysis. Random subsamples of 68 neurons from monkey D selected from the larger population of 124 to match those available in monkey H. Data were then smoothed and seqNMF was conducted using standard parameters. This procedure was repeated 100 times. The distributions of identified factors were compared to the full sample as well as the distribution of seqNMF runs returning 1 or 2 sequences (**Figure S9**).

## ACKNOWLEDGEMENTS

This work was supported by a National Institute of Mental Health BRAINS award to PHR (R01MH118638), a young investigator grant from the Brain and Behavior Foundation (NARSAD) to PHR, as well as NSF (NSF1926800) and NIMH (R01EB028166) grants to KR. We thank Dr Emily Mackevicius for advice on data analysis and Drs Brian Russ, Frederic Stoll, Matthew Perich, and Christian Marton for comments on earlier versions of the manuscript.

## REFERENCES

Adolphs, R., Damasio, H., Tranel, D., & Damasio, A.R. (1996) Cortical systems for the recognition of emotion in facial expressions. J Neurosci, 16, 7678–7687.

Adolphs, R., Tranel, D., Damasio, H., & Damasio, A.R. (1995) Fear and the human amygdala. J Neurosci, 15, 5879–5891.

Aggleton, J.P., Wright, N.F., Rosene, D.L., & Saunders, R.C. (2015) Complementary Patterns of Direct Amygdala and Hippocampal Projections to the Macaque Prefrontal Cortex. Cereb. Cortex, 25, 4351–4373.

Alexander, L., Gaskin, P.L.R., Sawiak, S.J., Fryer, T.D., Hong, Y.T., Cockcroft, G.J., Clarke, H.F., & Roberts, A.C. (2019) Fractionating Blunted Reward Processing Characteristic of Anhedonia by Over-Activating Primate Subgenual Anterior Cingulate Cortex. Neuron, 101, 307–320.e6.

An, X., Bandler, R., Ongur, D., & Price, J.L. (1998) Prefrontal cortical projections to longitudinal columns in the midbrain periaqueductal gray in macaque monkeys. J Comp Neurol, 401, 455–479.

Azab, H. & Hayden, B.Y. (2018) Correlates of economic decisions in the dorsal and subgenual anterior cingulate cortices. Eur. J. Neurosci., 47, 979–993.

Barbas, H. & Blatt, G.J. (1995) Topographically specific hippocampal projections target functionally distinct prefrontal areas in the rhesus monkey. Hippocampus, 5, 511–533.

Cash-Padgett, T., Azab, H., Yoo, S.B.M., & Hayden, B.Y. (2018) Opposing pupil responses to offered and anticipated reward values. Anim Cogn, 21, 671–684.

Cho, Y.T., Ernst, M., & Fudge, J.L. (2013) Cortico-amygdala-striatal circuits are organized as hierarchical subsystems through the primate amygdala. J Neurosci, 33, 14017–14030.

Crowe, D.A., Averbeck, B.B., & Chafee, M.V. (2010) Rapid sequences of population activity patterns dynamically encode task-critical spatial information in parietal cortex. J. Neurosci., 30, 11640– 11653.

Dowd, E.C. & Barch, D.M. (2010) Anhedonia and emotional experience in schizophrenia: neural and behavioral indicators. Biological psychiatry, 67, 902–911.

Dowd, E.C. & Barch, D.M. (2012) Pavlovian reward prediction and receipt in schizophrenia: relationship to anhedonia. PLoS One, 7, e35622.

Dunlop, B.W., Rajendra, J.K., Craighead, W.E., Kelley, M.E., McGrath, C.L., Choi, K.S., Kinkead, B., Nemeroff, C.B., & Mayberg, H.S. (2017) Functional Connectivity of the Subcallosal Cingulate Cortex And Differential Outcomes to Treatment With Cognitive-Behavioral Therapy or Antidepressant Medication for Major Depressive Disorder. Am J Psychiatry, 174, 533–545.

Enel, P., Wallis, J.D., & Rich, E.L. (2020) Stable and dynamic representations of value in the prefrontal cortex. Elife, 9.

Gabbott, P.L. & Rolls, E.T. (2013) Increased neuronal firing in resting and sleep in areas of the macaque medial prefrontal cortex. The European journal of neuroscience, 37, 1737–1746.

Ghashghaei, H.T., Hilgetag, C.C., & Barbas, H. (2007) Sequence of information processing for emotions based on the anatomic dialogue between prefrontal cortex and amygdala. Neuroimage, 34, 905–923.

Haber, S.N., Kim, K.S., Mailly, P., & Calzavara, R. (2006) Reward-related cortical inputs define a large striatal region in primates that interface with associative cortical connections, providing a substrate for incentive-based learning. Journal of Neuroscience, 26, 8368–8376.

Hahnloser, R.H.R., Kozhevnikov, A.A., & Fee, M.S. (2002) An ultra-sparse code underlies the generation of neural sequences in a songbird. Nature, 419, 65–70.

Harrison, N.A., Brydon, L., Walker, C., Gray, M.A., Steptoe, A., & Critchley, H.D. (2009) Inflammation causes mood changes through alterations in subgenual cingulate activity and mesolimbic connectivity. Biol Psychiatry, 66, 407–414.

Harvey, C.D., Coen, P., & Tank, D.W. (2012) Choice-specific sequences in parietal cortex during a virtual-navigation decision task. Nature, 484, 62–68.

Hochberg, Y. & Benjamini, Y. (1990) More powerful procedures for multiple significance testing. Stat Med, 9, 811–818.

Jezzini, A., Bromberg-Martin, E.S., Trambaiolli, L.R., Haber, S.N., & Monosov, I.E. (2021) A prefrontal network integrates preferences for advance information about uncertain rewards and punishments. Neuron, 109, 2339–2352.e5.

Ji, D. & Wilson, M.A. (2007) Coordinated memory replay in the visual cortex and hippocampus during sleep. Nat Neurosci, 10, 100–107.

Keedwell, P.A., Andrew, C., Williams, S.C., Brammer, M.J., & Phillips, M.L. (2005) The neural correlates of anhedonia in major depressive disorder. Biological psychiatry, 58, 843–853.

Kimmel, D.L., Elsayed, G.F., Cunningham, J.P., & Newsome, W.T. (2020) Value and choice as separable and stable representations in orbitofrontal cortex. Nat Commun, 11, 3466.

Lindquist, K.A., Wager, T.D., Kober, H., Bliss-Moreau, E., & Barrett, L.F. (2012) The brain basis of emotion: a meta-analytic review. Behav Brain Sci, 35, 121–143.

MacDonald, C.J., Lepage, K.Q., Eden, U.T., & Eichenbaum, H. (2011) Hippocampal “time cells” bridge the gap in memory for discontiguous events. Neuron, 71, 737–749.

Mackevicius, E.L., Bahle, A.H., Williams, A.H., Gu, S., Denisenko, N.I., Goldman, M.S., & Fee, M.S. (2019) Unsupervised discovery of temporal sequences in high-dimensional datasets, with applications to neuroscience. Elife, 8.

Mayberg, H.S., Brannan, S.K., Tekell, J.L., Silva, J.A., Mahurin, R.K., McGinnis, S., & Jerabek, P.A. (2000) Regional metabolic effects of fluoxetine in major depression: serial changes and relationship to clinical response. Biological psychiatry, 48, 830–843.

Mayberg, H.S., Liotti, M., Brannan, S.K., McGinnis, S., Mahurin, R.K., Jerabek, P.A., Silva, J.A., Tekell, J.L., Martin, C.C., Lancaster, J.L., & Fox, P.T. (1999) Reciprocal limbic-cortical function and negative mood: converging PET findings in depression and normal sadness. The American journal of psychiatry, 156, 675–682.

Mello, G.B.M., Soares, S., & Paton, J.J. (2015) A scalable population code for time in the striatum. Curr. Biol., 25, 1113–1122.

Mitz, A.R., Chacko, R.V., Putnam, P.T., Rudebeck, P.H., & Murray, E.A. (2017) Using pupil size and heart rate to infer affective states during behavioral neurophysiology and neuropsychology experiments. J Neurosci Methods, 279, 1–12.

Monosov, I.E. & Hikosaka, O. (2012) Regionally distinct processing of rewards and punishments by the primate ventromedial prefrontal cortex. The Journal of neuroscience: the official journal of the Society for Neuroscience, 32, 10318–10330.

Ongur, D., An, X., & Price, J.L. (1998) Prefrontal cortical projections to the hypothalamus in macaque monkeys. Journal of Comparative Neurology, 401, 480–505.

Pastalkova, E., Itskov, V., Amarasingham, A., & Buzsaki, G. (2008) Internally Generated Cell Assembly Sequences in the Rat Hippocampus. Science, 321, 1322–1327.

Paton, J.J., Belova, M.A., Morrison, S.E., & Salzman, C.D. (2006) The primate amygdala represents the positive and negative value of visual stimuli during learning. Nature, 439, 865–870.

Perich, M.G. & Rajan, K. (2020) Rethinking brain-wide interactions through multi-region “network of networks” models. Curr Opin Neurobiol, 65, 146–151.

Price, J.L. & Drevets, W.C. (2012) Neural circuits underlying the pathophysiology of mood disorders. Trends in cognitive sciences, 16, 61–71.

Prut, Y., Vaadia, E., Bergman, H., Haalman, I., Slovin, H., & Abeles, M. (1998) Spatiotemporal structure of cortical activity: properties and behavioral relevance. J Neurophysiol, 79, 2857–2874.

Rajan, K., Harvey, C.D., & Tank, D.W. (2016) Recurrent Network Models of Sequence Generation and Memory. Neuron, 90, 128–142.

Reitich-Stolero, T. & Paz, R. (2019) Affective memory rehearsal with temporal sequences in amygdala neurons. Nat Neurosci, 22, 2050–2059.

Rudebeck, P.H., Putnam, P.T., Daniels, T.E., Yang, T., Mitz, A.R., Rhodes, S.E., & Murray, E.A. (2014) A role for primate subgenual cingulate cortex in sustaining autonomic arousal. Proceedings of the National Academy of Sciences of the United States of America, 111, 5391–5396.

Schultz, W., Apicella, P., Ljungberg, T., Romo, R., & Scarnati, E. (1993) Reward-related activity in the monkey striatum and substantia nigra. Prog Brain Res, 99, 227–235.

Siegel, E.H., Sands, M.K., Van den Noortgate, W., Condon, P., Chang, Y., Dy, J., Quigley, K.S., & Barrett, L.F. (2018) Emotion fingerprints or emotion populations? A meta-analytic investigation of autonomic features of emotion categories. Psychol Bull, 144, 343–393.

Siegle, G.J., Thompson, W.K., Collier, A., Berman, S.R., Feldmiller, J., Thase, M.E., & Friedman, E.S. (2012) Toward clinically useful neuroimaging in depression treatment: prognostic utility of subgenual cingulate activity for determining depression outcome in cognitive therapy across studies, scanners, and patient characteristics. Arch. Gen. Psychiatry, 69, 913–924.

Smaragdis, P. (2006) Convolutive speech bases and their application to supervised speech separation. *IEEE Transactions on Audio*, Speech, and Language Processing, 15, 1–12.

Strait, C.E., Sleezer, B.J., Blanchard, T.C., Azab, H., Castagno, M.D., & Hayden, B.Y. (2016) Neuronal selectivity for spatial positions of offers and choices in five reward regions. J. Neurophysiol., 115, 1098–1111.

Taxidis, J., Pnevmatikakis, E.A., Dorian, C.C., Mylavarapu, A.L., Arora, J.S., Samadian, K.D., Hoffberg, E.A., & Golshani, P. (2020) Differential Emergence and Stability of Sensory and Temporal Representations in Context-Specific Hippocampal Sequences. Neuron, 108, 984–998.e9.

Vaz, A.P., Wittig, J.H., Inati, S.K., & Zaghloul, K.A. (2020) Replay of cortical spiking sequences during human memory retrieval. Science, 367, 1131–1134.

Wallis, C.U., Cockcroft, G.J., Cardinal, R.N., Roberts, A.C., & Clarke, H.F. (2019) Hippocampal Interaction With Area 25, but not Area 32, Regulates Marmoset Approach-Avoidance Behavior. Cereb. Cortex, 29, 4818–4830.

Zeredo, J.L., Quah, S.K.L., Wallis, C.U., Alexander, L., Cockcroft, G.J., Santangelo, A.M., Xia, J., Shiba, Y., Dalley, J.W., Cardinal, R.N., Roberts, A.C., & Clarke, H.F. (2019) Glutamate Within the Marmoset Anterior Hippocampus Interacts with Area 25 to Regulate the Behavioral and Cardiovascular Correlates of High-Trait Anxiety. J. Neurosci., 39, 3094–3107.

