## Supplemental information for "Temporally specific patterns of neural activity in interconnected corticolimbic structures during reward anticipation"

### SUPPLEMENTARY INFORMATION

**Title: Temporally specific sequences of neural activity across interconnected corticolimbic structures during reward anticipation**

**Authors**: Megan E. Young*+, Camille Spencer-Salmon*, Clayton P. Mosher, Sarita Tamang, Kanaka Rajan, and Peter H. Rudebeck

**ADDRESS**:
Icahn School of Medicine at Mount Sinai, One Gustave L. Levy Place, New York, NY, 10029, USA

* These authors contributed equally to this work
[+] Current address: Department of Integrative Neuroscience, University of Toyama, Toyama, 930-8555, Japan

**Figures:** 9
**Tables:** 0

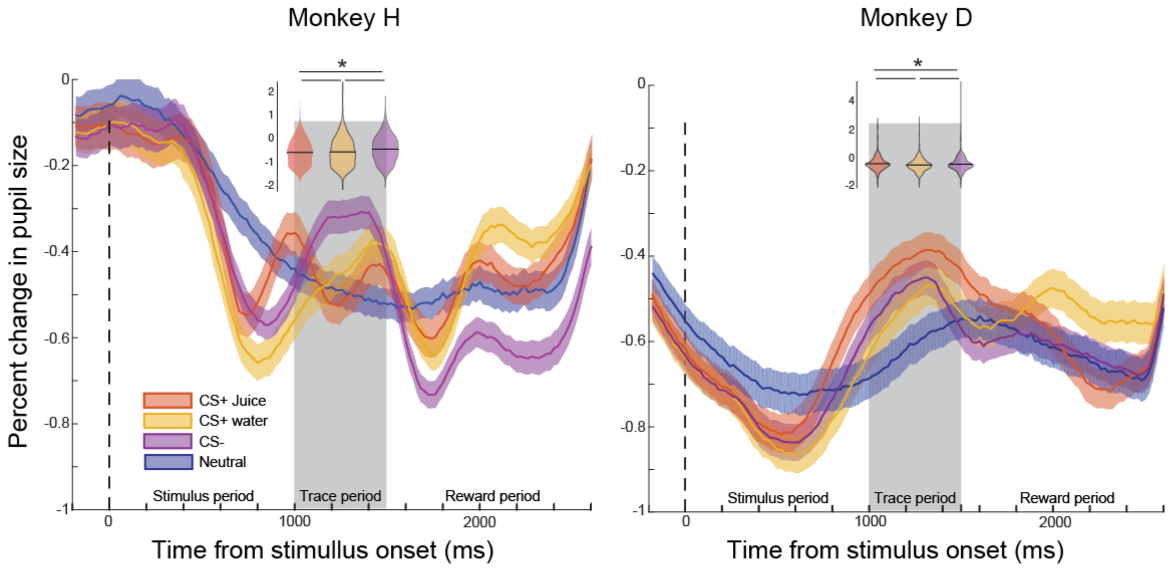


**Supplemental Figure S1: Pupil size responses to different trial types.** Corresponds to Figure 1. Pupil size as a function of time (percent change, mean ± SEM) for monkeys H (left) and D (right) on CS+$^{juice}$ (red), CS+$^{water}$ (yellow), or CS– trials (purple). Shaded regions or error bars show SEM. Stimulus onset, offset and time of reward delivery are marked by dotted lines. The filled gray bar marks the period on which the inset violin plots of mean and distribution of conditions where conditioned stimuli were presented are based. Analyzing the mean pupil size within this period of time across the different sessions revealed that both monkeys exhibited statistically significant responses to the different conditions (monkey H: $F(3,5133)=12$; $p<0.0001$; monkey D: $F(3,3880)=14.76$; $p<0.0001$), albeit with different patterns (pairwise comparisons, $p<0.05$, monkey H, CS- > CS+$^{water}$ > CS+$^{juice}$; monkey D, CS+$^{juice}$ > CS-> CS+$^{water}$ ). Horizontal lines on the violin plot mark significant comparisons at $p < 0.05$.

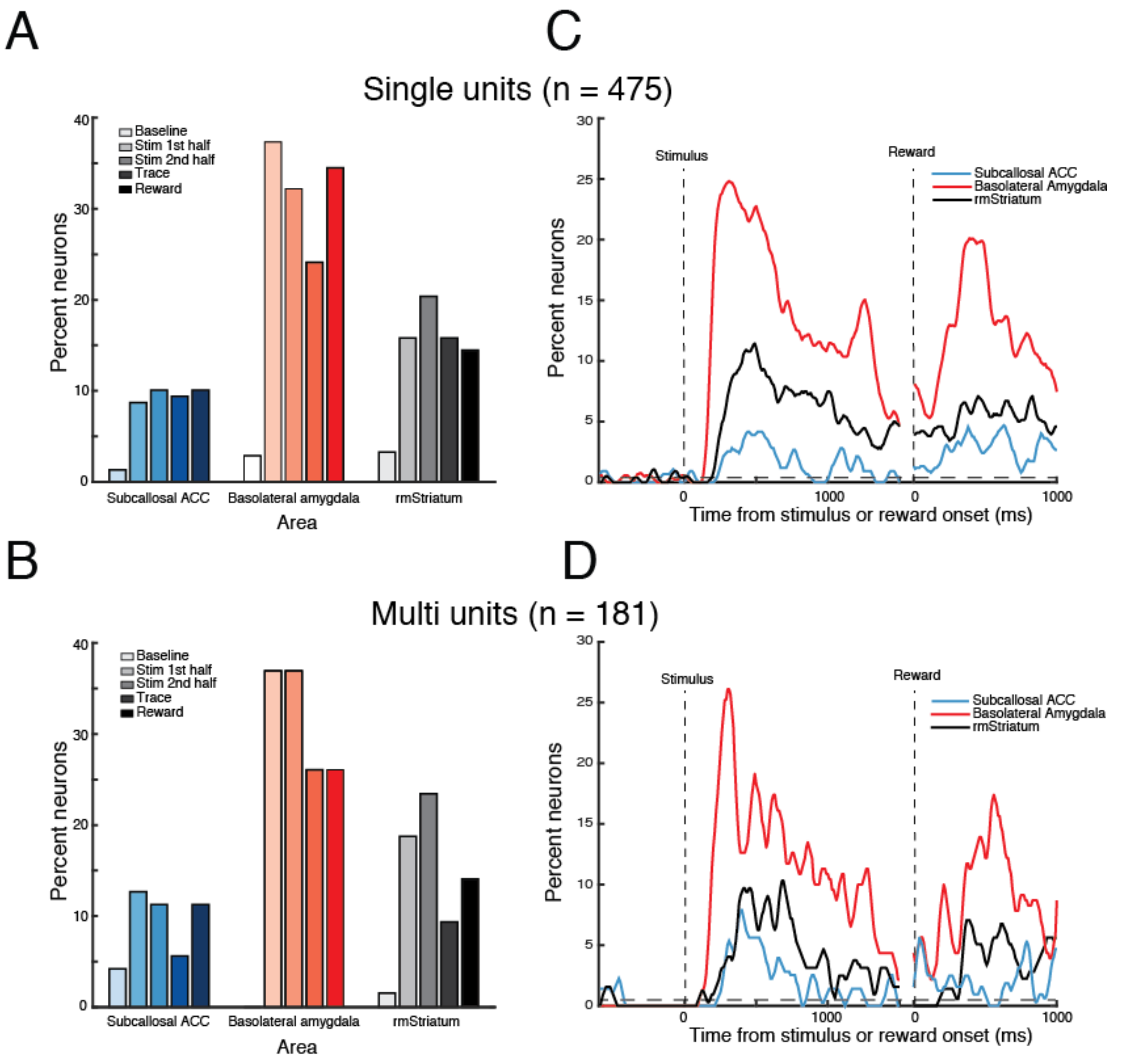


**Supplemental Figure S2: Single neuron and multi-unit encoding of trial type in the Pavlovian task.** Corresponding to Figure 2. **A/B**) Percent of single neurons (**A**) and multi-unit recordings (**B**) in subcallosal ACC, BLA and rostromedial striatum classified by a sliding ANOVA as encoding a the different task conditions during either the *baseline period* (1.0 s before stimulus onset), the *stimulus period 1st half* (0–0.5 s after stimulus onset), the *stimulus period 2nd half* (0.5–1 s after stimulus onset), or the *reward period* (0-0.5 s after reward onset). **C/D**) Time course of single neuron (**C**) and multi-unit (**D**) stimulus–reward encoding in subcallosal ACC (blue), BLA (red) and rostromedial striatum (black) following the presentation of the conditioned stimuli. Vertical dashed lines correspond to the onset of the stimulus (left) and reward (right). Horizontal dashed line depicts the data derived false discovery rate at each timepoint.

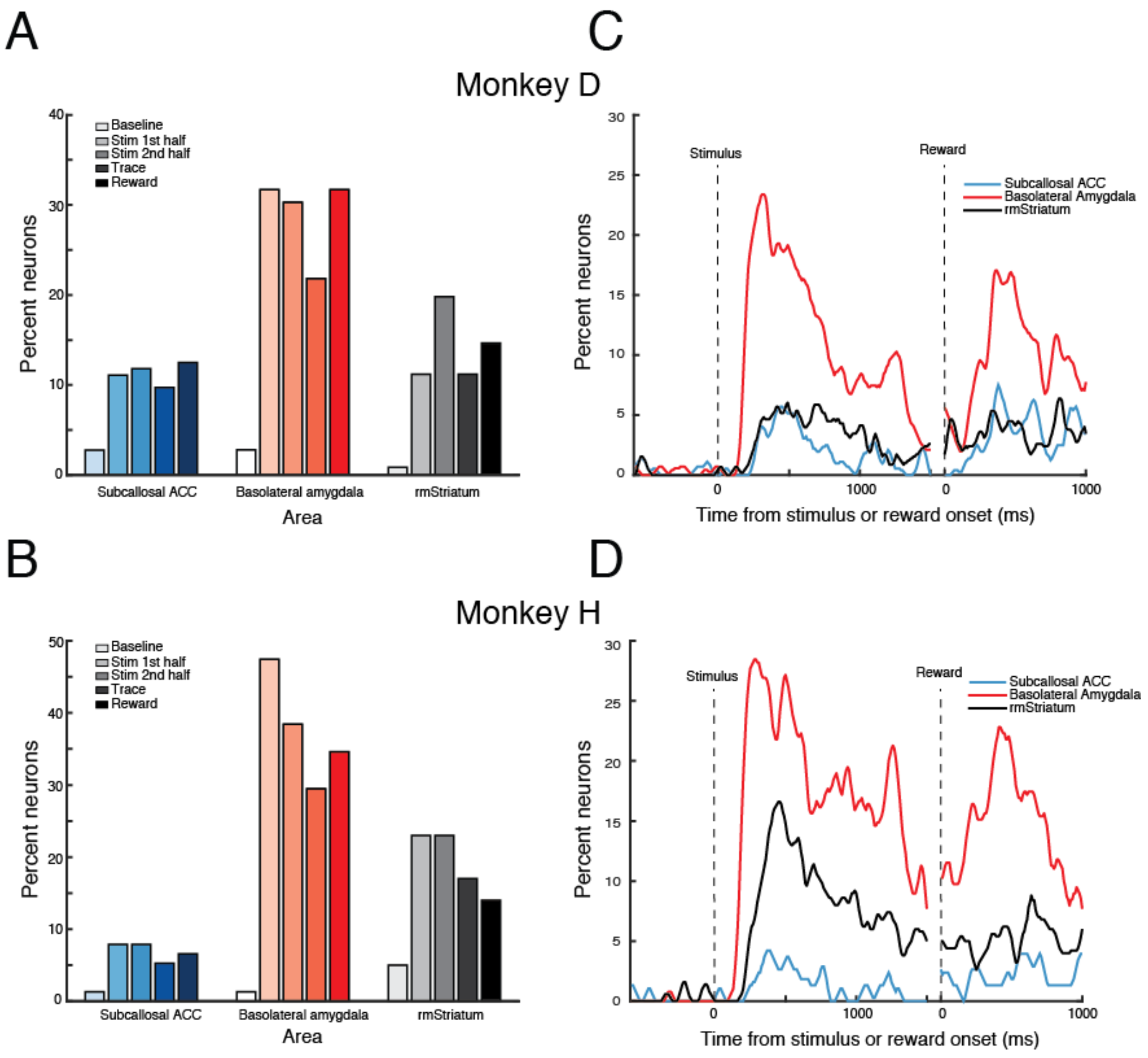


**Supplemental Figure S3**: **Neural activity in subcallosal ACC, BLA and rostromedial striatum during the Pavlovian task by subject.** Corresponds to Figure 2. Single neuron encoding of trial type in the Pavlovian task for monkey D and H (Corresponding to Figure 2). **A/B**) Percent of single neurons in monkey D (**A**) and monkey H (**B**) in subcallosal ACC, BLA and rostromedial striatum classified by a sliding ANOVA as encoding a the different task conditions during either the *baseline period* (0.5 s before the onset of stimulus), the *stimulus period 1$^{st}$ half* (0–0.5 s after the onset of the stimulus), *stimulus period 2$^{nd}$ half* ( 0.5–1 s after the onset of stimulus), the *trace interval* (1-1.5s after the onset of the stimulus), or the *reward period* (0-0.5s after reward onset). **C/D**) Time course of single neuron stimulus–reward encoding for Monkey D (**C**) and Monkey H (**D**) in subcallosal ACC (blue), BLA (red) and rostromedial striatum (black) following the presentation of the conditioned stimuli. Vertical dashed lines correspond to the onset of the stimulus (left) and reward (right).

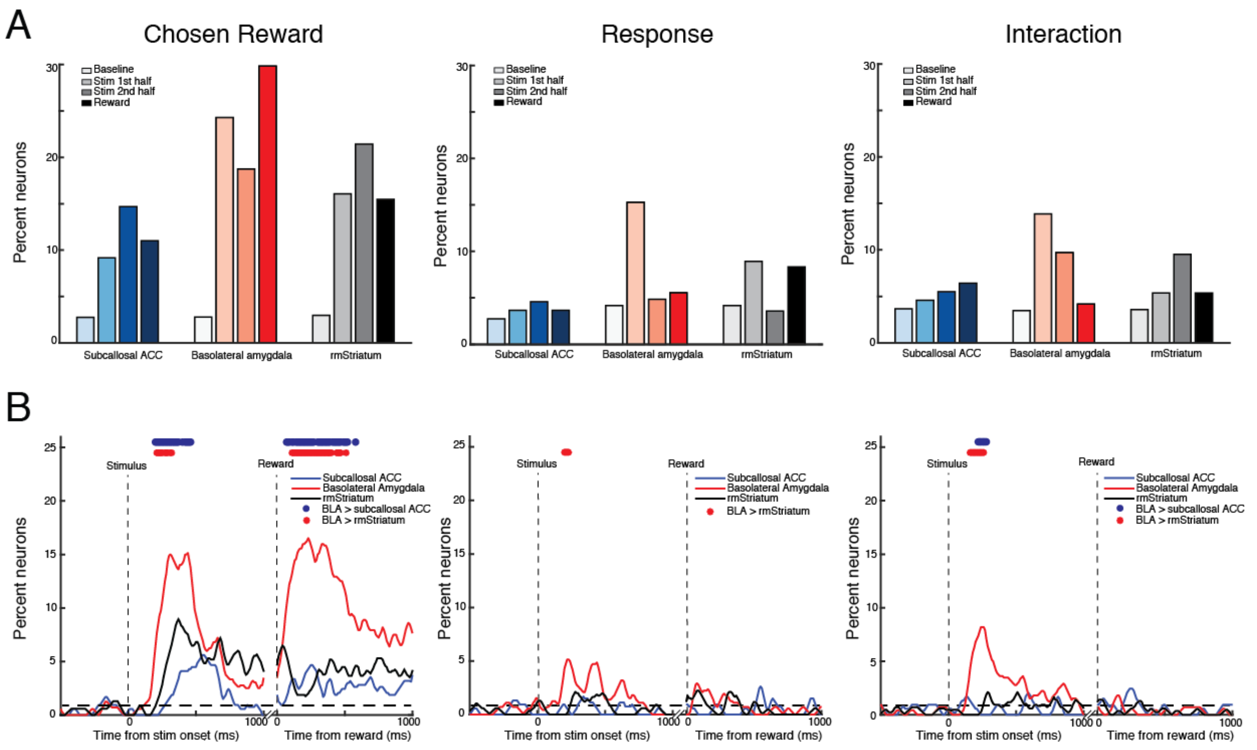


**Supplemental Figure S4**: **Neural activity in subcallosal ACC, BLA and rostromedial striatum during the instrumental task.** Corresponds to Figure 4. **A**) Percent of neurons in subcallosal ACC (shaded blue), BLA (shaded red) and rostromedial striatum (shaded grayscale) classified by a sliding ANOVA as encoding chosen reward (right), movement direction (middle) and interaction between reward and movement direction (right) during either the *baseline period* (0.5 s before the onset of the stimuli), the *stimulus period 1$^{st}$ half* (0–0.5 s after the onset of the stimulus), the *stimulus period 2$^{nd}$ half* ( 0.5–1 s after the onset of the stimulus) or *reward period* (0-0.5s after reward onset). **B**) Time course of encoding of chosen reward (left), movement direction (middle) and interaction between chosen reward and movement direction (right) in subcallosal ACC (blue), BLA (red) and rostromedial striatum (black) following stimulus or reward onset. Red and blue dots at the top indicate significant differences in the proportion of neurons between areas ($p<0.0167$, Gaussian approximation test with false discovery rate correction). Rostromedial striatum and subcallosal ACC comparison not shown as all p values >0.05. Vertical dashed lines correspond to the onset of stimuli and reward delivery. Horizontal dashed line depicts the data derived false discovery rate at each timepoint.

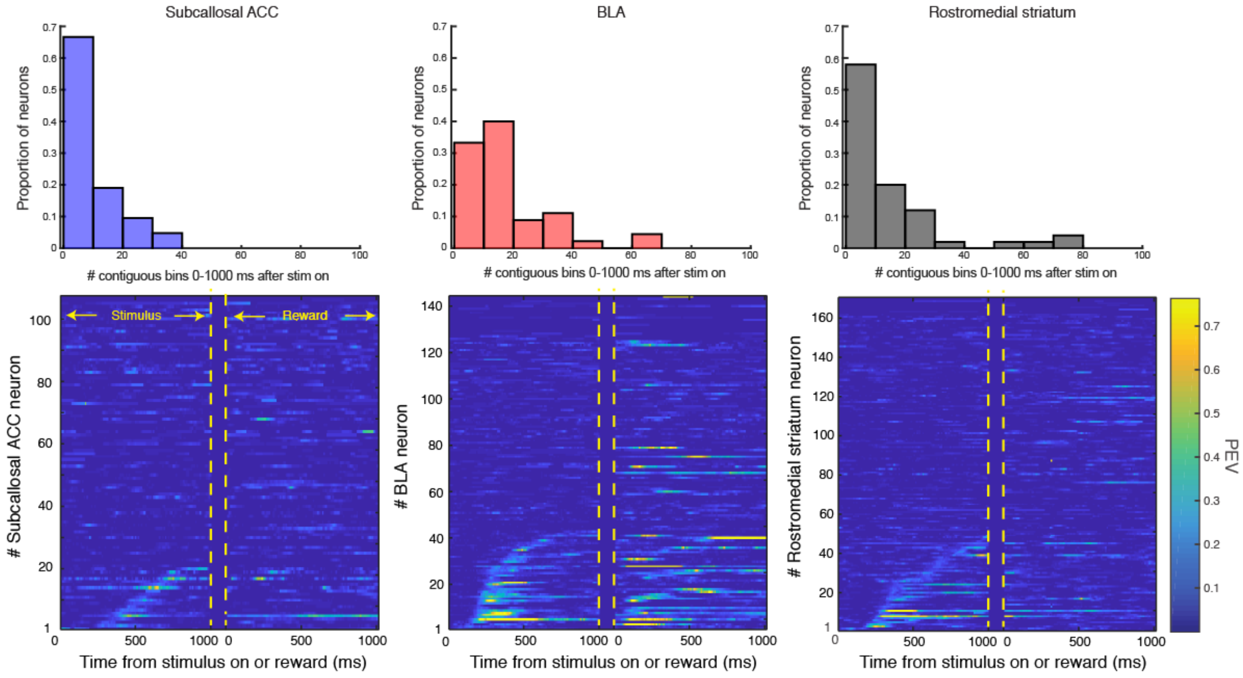


**Supplemental Figure S5: Timing and length of encoding of chosen reward in subcallosal ACC, BLA and rostromedial striatum in the instrumental task.** Corresponds to **Figures 3** and **4**. Percent explained variance (PEV) associated with the different chosen reward for each neuron (bottom). In the plots of PEV, neurons are sorted according to when they were classified as being significantly modulated by chosen reward. Lighter or 'hotter' colors are associated with higher explained variance. Because of the variable response time of the monkeys, data are temporally realigned for the reward period and there is a period which has been intentionally left blank/dark blue between 1000-1200 ms. Histograms (top) show the amount of time encoding for subcallosal ACC (left), BLA (middle), rostromedial striatum (right) relative to stimulus and reward onset. Time encoding is based on the number of contiguous bins (10 ms step) where each neuron was classified as encoding. Kruskall-Wallis test $\chi^2>6.24$, $p<0.05$, BLA versus subcallosal ACC or rostromedial striatum, $\chi^2>5.66$, $p<0.05$. No other pairwise comparisons reached the threshold for statistical significance.

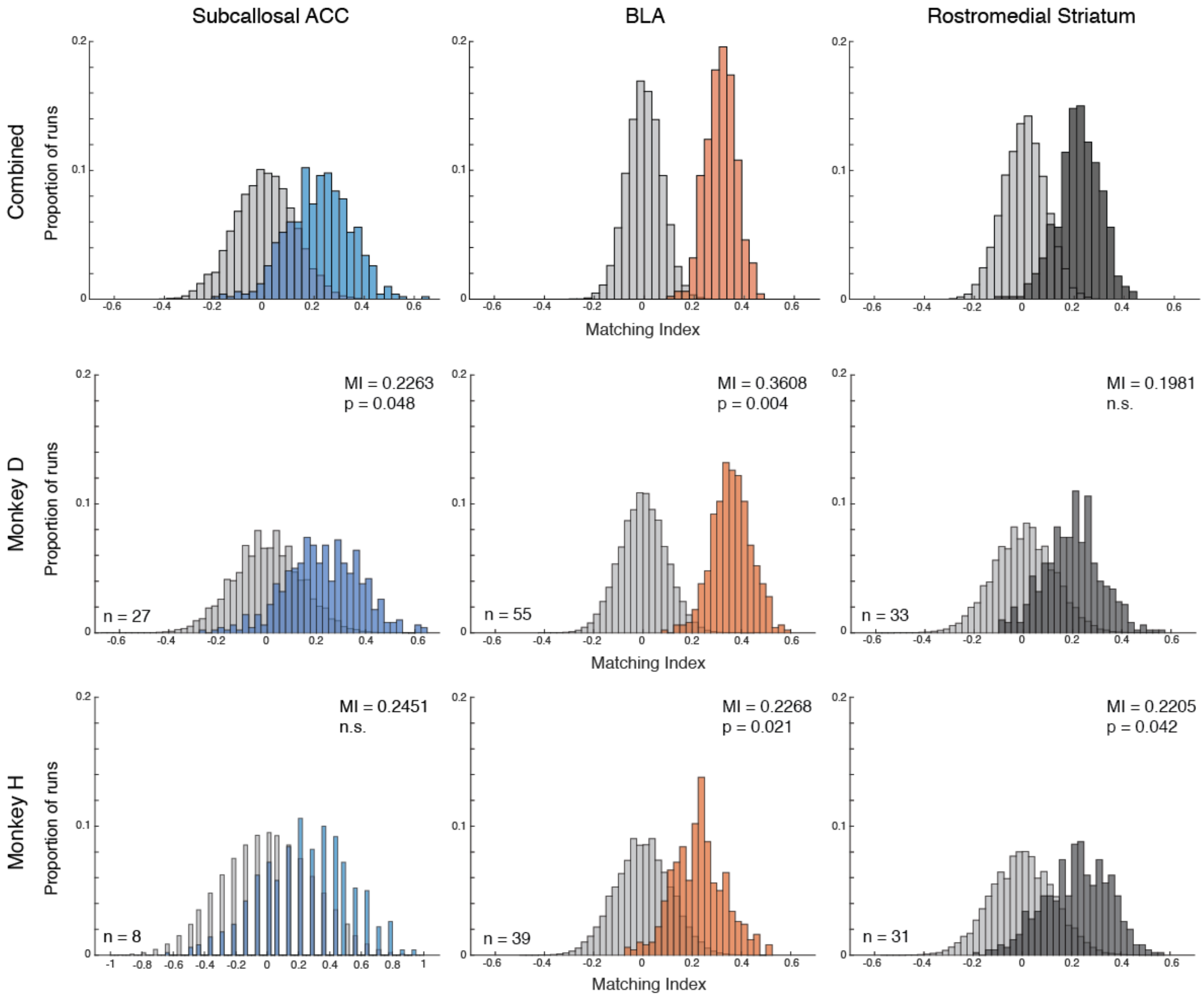


**Supplementary Figure S6: Matching index in subcallosal ACC, BLA and rostromedial striatum for combined, and monkeys D and H.** Corresponds to **Figure 5C**. Distribution of matching indexes for subcallosal ACC (left), BLA (middle), and rostromedial striatum (right) across the 500 runs for combined (top), monkey D (middle) and monkey H (bottom). Shuffled data are in grey and actual data are in corresponding colors.

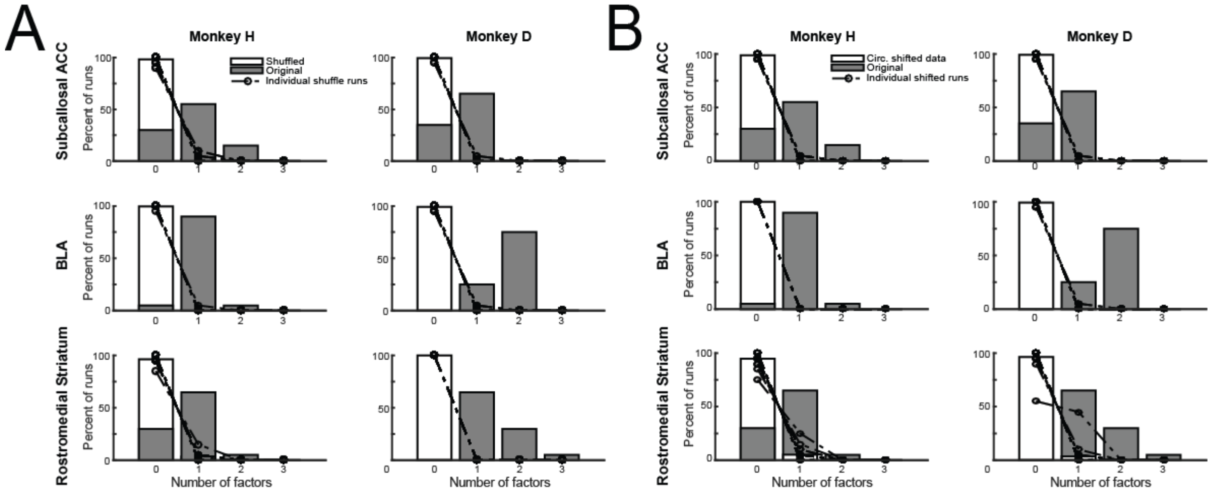


**Supplemental Figure S7: seqNMF spike-time shuffle circular shuffle control analyses.** Corresponds to Figures 7D, 8A and 8. **A)** Histograms of the number of significant factors identified from the 20 training and testing runs of seqNMF for monkeys H and D for subcallosal ACC (top), BLA (middle) and rostromedial striatum (bottom). **B**) Histograms of the number of significant factors identified from the 20 training and testing runs of seqNMF for monkeys H and D for subcallosal ACC (top), BLA (middle) and rostromedial striatum (bottom). In both **A** and **B**, the distribution of factors identified from original unshifted data from the CS/trace period are shown as gray bars. The mean number of factors identified from temporally circularly shifted data are shown as unfilled bars. Data were randomly shifted within each trial segment and seqNMF was run. This process was repeated 20 times and dotted lines represent the factors recovered on individual runs. Individual runs are often obscured as they overlap.

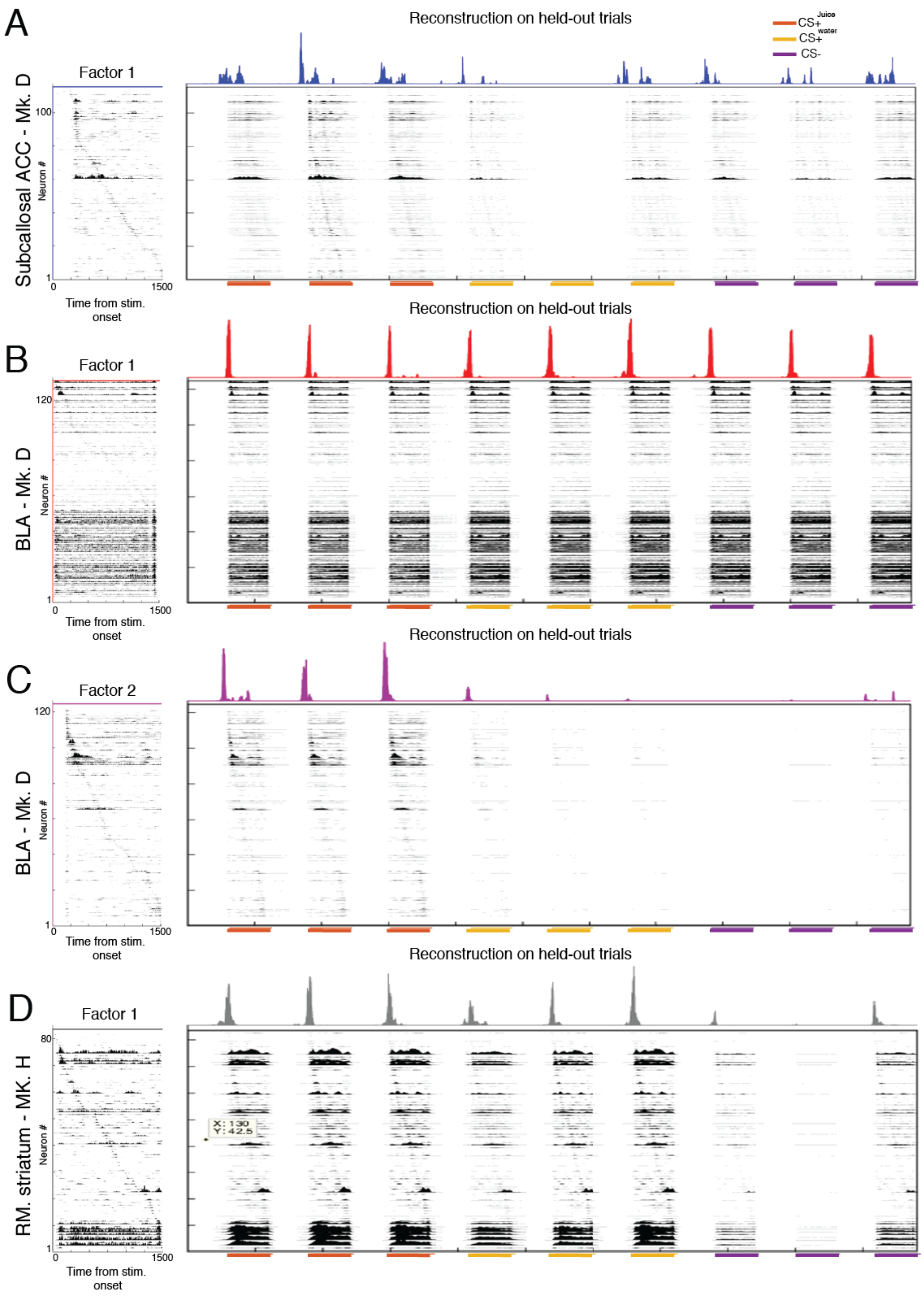


**Supplemental Figure S8: Reconstruction of significant factors on held out trials in subcallosal ACC, BLA and rostromedial striatum.** Corresponds to Figures 7 and 8. **A, B, C, D**)

Significant extracted factors (left) and reconstructions of those factors on held-out trials from three CS+$^{juice}$, CS+$^{water}$, and CS- trials (right) from pseudo-ensembles of subcallosal ACC (A), BLA (B,C) and rostromedial striatum neurons (D). Within each factor neurons are sorted according to the time point of their peak activation. The period shown is from 0 to 1500 ms after stimulus onset which includes both stimulus and trace periods. In the reconstructions, colored plots (top) on each trial show the temporal loadings over time for CS+$^{juice}$, CS+$^{water}$, and CS-. The average of these temporal loadings from each trial type is used to make the bar plots shown in **Figures 7G, H** and **8 E, F**.

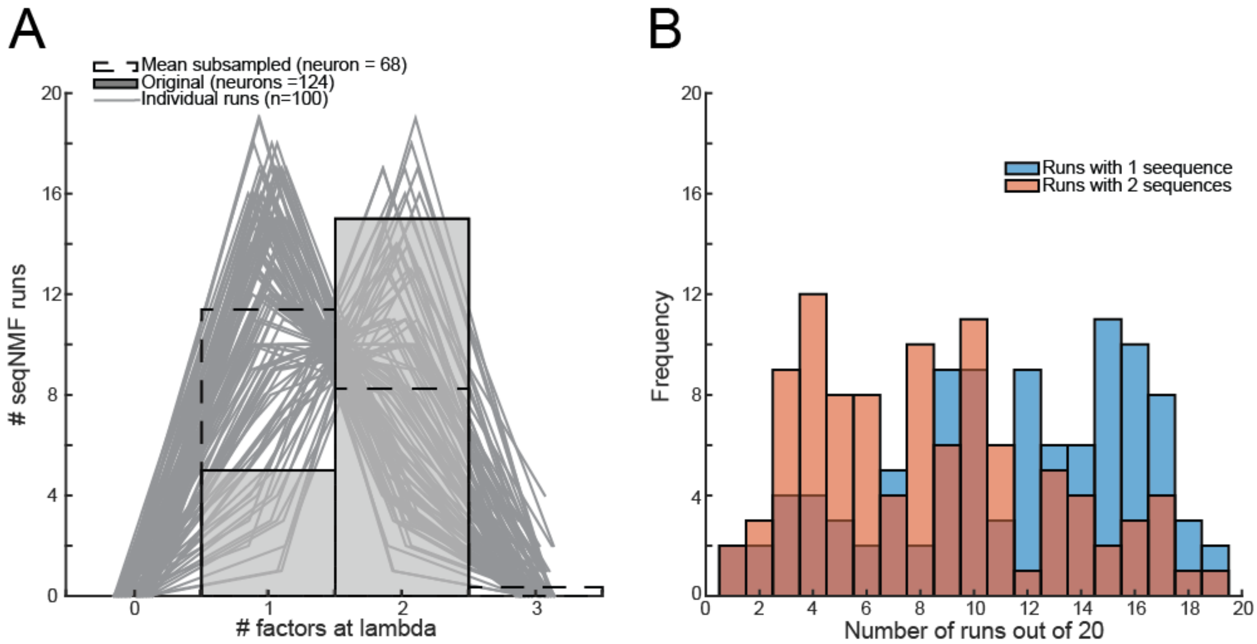


**Supplemental Figure 9: seqNMF subsample analysis on Monkey D amygdala.** Corresponds to Figure 8F. A) Mean number of sequences identified in subsampled (dashed line) or original data (grey bars). For the subsample analysis, pseudoensembles of 68 neurons from BLA of monkey D were randomly selected from the total pool of 124 and seqNMF was run on the data using parameters applied to the original dataset. Using this approach, 20 runs of seqNMF were performed to determine the number of sequences identified. This process was repeated 100 times, yielding 2,000 runs in total. The number of sequences identified in the data from the 20 training runs from each of the 100 iterations is plotted as individual lines. B) Histogram of the frequency of seqNMF iterations where either 1 (blue) or 2 (red) sequences were returned. Statistical comparison of the frequency revealed a that seqNMF was more likely to return a single rather than two sequences when only 68 neurons were included in the analysis (Kruskal-Wallis test, $F(1,199)=21.45$, $p<0.001$).
